# Non-DNA-damaging DNA-PK activation improving hearing and prolonging life due to NAD^+^ and SIRT upregulation

**DOI:** 10.1101/2025.04.18.649305

**Authors:** Yohei Honkura, Takehiro Suzuki, Ryota Kujirai, Shinichiro Kanno, Yotaro Matsumoto, Yoshikazu Tanaka, Hengphasatporn Kowit, Satoru Nagatoishi, Alexander Tyshkovskiy, Tomoko Kasahara, Yoshiyasu Tongu, Zhang Bohan, Hitomi Kashiwagi, Chika Saegusa, Msato Fujioka, Reina Usami, Shunsuke Chikuma, Yukiko Tokifuji, Tetsuro Matsuhashi, Yoshitsugu Oikawa, Hiroka Komatsu, Kei Murayama, Takehito Sugasawa, Fumika Nanto-Hara, Kensei Taguchi, Daisuke Saigusa, Chitose Suzuki, Takeya Sato, Jun Suzuki, Yuji Owada, Kuniyasu Niizuma, Hidenori Endo, Ko Hashimoto, Takafumi Toyohara, Kohei Tsumoto, Paul Anderson, Vadim N. Gladyshev, Ken-ichiro Hayashi, Yukio Katori, Yoshihisa Tomioka, Takaaki Abe

**Affiliations:** Department of Otolaryngeal Surgery, Tohoku University Graduate School of Medicine, Sendai, Japan; Division of Medical Science, Tohoku University Graduate School of Biomedical Engineering, Sendai, Japan; Department of Nephrology, Tohoku University Graduate School of Medicine, Sendai, Japan; Laboratory of Oncology, Pharmacy Practice and Sciences, Tohoku University Graduate School of Pharmaceutical Sciences, Sendai, Japan; Department of Clinical Biology and Hormonal Regulation, Tohoku University Graduate School of Medicine, Sendai, Japan; Department of Biomolecular Sciences, Tohoku University Graduate School of Life Sciences, Sendai, Japan; Center for Computational Sciences, University of Tsukuba, Japan; Institute of Medical Sciences, The University of Tokyo, Tokyo, Japan; Division of Genetics, Department of Medicine, Brigham and Women’s Hospital, Harvard Medical School, Boston, MA, USA; Department of Molecular Genetics, Kitasato University School of Medicine, Tokyo, Japan; Department of Microbiology and Immunology, Keio University School of Medicine, Tokyo, Japan; Institute of Biotechnology, College of Life Sciences and Medicine National Tsing-Hua University, Hsinchu, Taiwan; Department of Pediatrics, Tohoku University Graduate School of Medicine, Sendai, Japan; Department of Hematology, Tohoku University Graduate School of Medicine, Sendai, Japan; Diagnostics and Therapeutics of Intractable Disease, Intractable Disease Research Center and Department of Pediatrics, Juntendo University Faculty of Medicine, Tokyo, Japan; Division of Sports Medicine, University of Tsukuba, Tsukuba, Japan; Division of Nephrology, Department of Medicine, Kurume University School of Medicine, Kurume, Japan; Laboratory of Biomedical and Analytical Sciences, Faculty of Pharma-Sciences, Teikyo University, Japan; Department of Anatomy, Tohoku University Graduate School of Medicine, Sendai, Japan; Department of Neurosurgical Engineering, Graduate School of Biomedical Engineering, Tohoku University, Sendai, Japan; Department of Translational Neuroscience, Tohoku University Graduate School of Medicine, Sendai, Japan; Department of Neurosurgery, Tohoku University Graduate School of Medicine, Sendai, Japan; Department of Orthopaedic Surgery, Tohoku University Graduate School of Medicine, Sendai, Japan; Division of Rheumatology, Inflammation, and Immunity at Brigham and Women’s Hospital, Harvard Medical School, Boston, MA, USA; Department of Bioscience, Okayama University of Science, Okayama, Japan

**Author notes:** **To whom correspondence should be addressed:** Takaaki Abe, M.D., Ph.D., Division of Medical Science, Tohoku University Graduate School of Biomedical Engineering, Sendai 980-8574, Japan. these authors contributed equally to this work (Yohei Honkura, Takehiro Suzuki, Ryota Kujirai, Shinichiro Kanno). Division of Meat Animal and Poultry Research, Institute of Livestock and Grassland Science, National Agriculture and Food Research Organization (NILGS), Tsukuba, Japan (Fumika Nanto-Hara).

**Keywords:** hearing loss, sirtuin, ATP, NAD^+^, NAMPT, TRIM28, sirtuin, PARP-1, DNA-PK

## Abstract

Emerging evidence strongly supports a close relationship between age-related hearing loss and frailty, highlighting the importance of early detection and intervention. Recently, we invented a mitochondria-homing drug named mitochonic acid 5 (MA-5), that increases the adenosine triphosphate (ATP) levels, rescue mitochondrial function, and protect tissue damages. Currently, the phase I clinical trial has been finished in Japan (jRCT2031210495) and the phase 2 clinical trial has already been approved by PMDA.

Here we show that MA-5 improved various types of hearing loss in mouse models. Structural chemical bioanalysis revealed that MA-5 is a mixture of equal amount of S- and R- enantiomer and both S- and R- enantiomer increase ATP by binding mitochondrial protein, mitofilin. However, S-enantiomer significantly increased the NAD^+^ levels by binding to the NAD^+^-producing key enzyme nicotinamide phosphoribosyltransferase (NAMPT). Moreover, the S-enantiomer increased the sirtuin 1 protein by suppressing polyubiquitination induced by tripartite motif containing 28 (TRIM28) phosphorylation which was triggered by DNA-dependent protein kinase (DNA-PK) activation in the absence of DNA damage. Transcriptomic signatures showed that the signature of MA-5 shows an inverse correlation with aging and mortality and is oriented in the same direction as the OSKM-related iPSCs, suggesting the modification of aging pathways. Oral administration of MA-5 to mitochondrial disease model mouse showed increased survival. Our findings suggest that, in addition to enhancing ATP levels, the coordinated regulation of NAD^+^ metabolism, SIRT protein expression, and DNA-PK activity-constituting a novel therapeutic triad may contribute to the amelioration of hearing impairment and mitochondrial dysfunction, thereby improving life prognosis.

## Introduction

Hearing loss is associated with higher rates of frailty, hospitalization, dementia and depression as well as reduced quality of life^1^. The primary causes of hearing loss are age-related hearing loss (ARHL), noise-induced hearing loss (NIHL), and drug-induced hearing loss (DRHL)^2^. Mitochondria are key organelles that regulate a variety of cellular processes, including energy and free radical generation, apoptosis, and the modulation of hearing. Mitochondrial dysfunction causes ARHL^3^, NIHL^4^, and DRHL^5^. In addition, mutations in mitochondrial genes are a major cause of genetic hearing loss^2^. Many studies have suggested that not only mitochondrial dysfunction but also reduced nicotinamide adenine dinucleotide (NAD^+^) levels and loss of the expression level of sirtuins (SIRTs) are major causes of hearing loss^6^. Previous studies showed that SIRT1 and SIRT3 are involved in the pathogenesis of ARHL^7^, NIHL^8^, and DRHL^9^. These findings indicate that controlling mitochondrial damage with increasing ATP and NAD^+^, as well as up-regulating the SIRTs expression level are pivotal strategies for overcoming hearing loss. Recently, we invented a mitochondria-homing drug named mitochonic acid 5 (MA-5)^10–12^, that increases the adenosine triphosphate (ATP) levels, rescue mitochondrial disease fibroblasts’ death, and protect tissue damages in a mitochondrial DNA-defected disease model called “MitoMouse”^10^. MA-5 is now under phase 2 clinical trial. MA-5 is a synthetic indole derivative with a racemic mixture of R- and S-enantiomers^11^ and MA-5 has been developed as a drug for the treatment of rare diseases based on its pharmacological properties. However, in general, it would be desirable to develop the enantiomeric form when developing a racemic compound as a drug because each enantiomer exhibits pharmacological activities that can be null, similar, different, or opposite. For example, the use of thalidomide during gestation causes multiple birth defects and malformations; these side effects are solely attributed to the enantiomer. Here, we showed that the S-enantiomer of MA-5 increased not only the ATP but also NAD^+^ levels by binding to NAD^+^-producing key enzyme nicotinamide phosphoribosyltransferase (NAMPT). This effect was weak for the R-enantiomer and was considered unique to the S-enantiomer. Moreover, the S-enantiomer increased the SIRT1–7 protein levels by promoting the non-DNA-damaging phosphorylation of tripartite motif containing 28 (TRIM28) following the suppression of the polyubiquitination of SIRT proteins. This TRIM28 phosphorylation by S-enantiomer occurs independently of ataxia-telangiectasia-mutated (ATM) or ATM- and Rad3-related (ATR) phosphorylation, but may be up-regulated to DNA-dependent protein kinase (DNA-PK). Although it is believed that activation of DNA-PK occurs during DNA damage (e.g. aging, radiation anti-cancer drugs, etc.,) there are a few reports in the DNA-PK activation mechanism in the absence of DNA damage. In this study, we elucidated the multifaceted mechanisms by which the drug enhances ATP, NAD^+^, and SIRT proteins expression, and demonstrated its beneficial effects on impaired healing and lifespan in mitochondrial-related diseases.

## RESULTS

### MA-5 improves mitochondria-related and various types of hearing loss

We previously reported that MA-5 is effective on ” MitoMouse”, which has a genetic mutation in the mitochondrial DNA^13^. However, the difficulty of introducing the mutant gene in to mitochondrial DNA and of obtaining diseased individuals and the heteroplasmy rate differences among tissues, prevent the large-scale animal studies. On the other hand, the NADH dehydrogenase (ubiquinone) Fe-S protein 4 (Ndufs4) knockout (KO) mouse carries a mutation in a nuclear gene, and the resulting reduction in complex I function leads to fatal encephalomyelopathy, resembling that observed in patients with Leigh syndrome^14^. Recently, we reported that Ndufs4 KO mice show progressive hearing loss assessed by the auditory brainstem response (ABR) test recognized as a mitochondrial disease-related hearing loss^15^. At P28, no significant difference in ABR hearing thresholds was observed in both wild-type (WT) and Ndufs4 KO mice (Extended Data Fig. 1a). Under the condition, we administered MA-5 to mice from postnatal day 28 (P28) to P62 and examined the auditory capacity (Fig. 1a). However, at P62, the ABR thresholds were higher than those of WT mice in all frequencies and MA-5 improved the hearing threshold of Ndufs4 KO mice at frequencies of 8, 12, 16, and 32 kHz (Fig. 1a, left). Hearing loss typically involves impairment of the hair cells of the organ of Corti, stria vascularis, and spiral ganglion cells (SGCs)^16^, and histological examination of cochlear revealed that MA-5 restored the reduced SGC densities in Ndufs4 KO mice (Fig. 1a, right). In addition, since hearing loss naturally occurs along with aging, we next examined whether MA-5 could improve hearing capacity after the onset of impairment. At P40, the hearing capacity of Ndufs4 KO mice was started to decrease compared with P28 (Extended Data Fig. 1b), and we started to administer MA-5 at P40 and examined the hearing function at P60. As a result, MA-5 led to improvements in hearing at 8, 12, and 16 kHz in Ndufs4 KO mice at P60 (Fig. 1b, left). Histological examination also demonstrated that MA-5 restored the SGC densities (Fig. 1b, right). These results further indicated that MA-5 improved hearing function even after the onset of hearing function decline, which is similar mechanism of age-related hearing loss (ARHL). Noise-induced hearing loss (NIHL) is also a common type of sensorineural hearing loss. The synapses between inner hair cells (IHCs) are the most vulnerable by large sound and mitochondrial dysfunction is associated with the damage^1^. Accordingly, we next examined the effects of MA-5 in NIHL mice. ABR threshold was measured after noise exposure for in wild type mice in all frequencies. After administration of MA-5 the threshold was decreased at 8, 12, and 16 kHz for 4 h (Fig. 1c, left) and all frequencies for 7 days (Fig. 1c, middle) in both 1 and 10 μg/kg of MA-5. Histological examination revealed that the reduced density of IHCs was ameliorated by MA-5 (Fig. 1c, right). This suggests that the use of MA-5 may be effective in preventing NIHL caused by environmental factors. Cisplatin is widely used in the treatment of various types of cancer, however, it causes severe ototoxicity with irreversible hearing loss. Cisplatin causes mitochondrial dysfunction in sensory hair cells, the SGC and the lateral wall of the cochlea, which are its primary targets^17^. Therefore, we next examined the effects of MA-5 on cisplatin-induced hearing loss. After 5 days of cisplatin treatment, the ABR threshold increased in all frequencies compared with the control group (Fig. 1d, left). Under the condition, the MA-5 exhibited a significant reduction in the ABR threshold elevation, with a mean reduction of ∼10 dB at frequencies of 4, 8, 12, and 16 kHz. Histological examination revealed that MA-5 restored the mean SGC density and hair cell survival and stria vascularis structure (Fig. 1d, right). To further evaluate the protective effect of MA-5 against cisplatin-induced hearing loss, cochlear explants were pre-treated with MA-5 for 1 h before cisplatin exposure, and the number of hair cells in the apical, middle, and basal turns was determined. Cisplatin damaged the cochlear IHCs and outer hair cells (OHCs), especially in the basal turn, in a dose-dependent manner (Fig. 1e, right). The damage was more severe in the OHC three lines compared with that in the IHCs. Under the condition, MA-5 prevented cisplatin-induced damage in the basal, middle, and apical turns (Fig. 1e, right). These data clearly demonstrated that MA-5 has the potential to be effective for all types of hearing loss, and its clinical application could be of great help in alleviating hearing loss.

**Figure 1.**
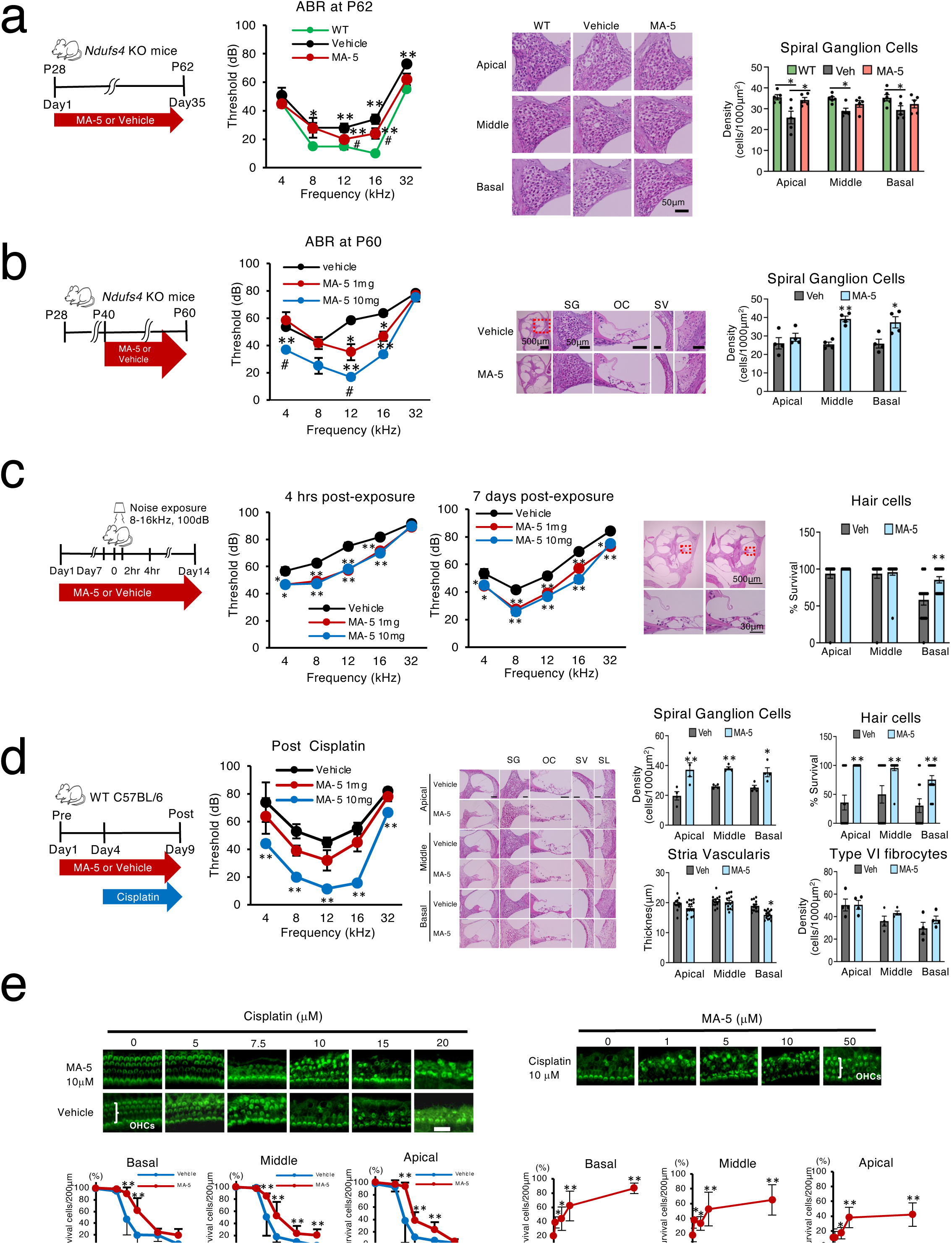
MA-5 ameliorates hearing deficits in four distinct models of hearing loss. **a,** Auditory brainstem response (ABR) test of WT and Ndufs4 KO mice administrated with Vehicle or MA-5 (1mg/kg) for 5 weeks from postnatal day 28 (P28) to P62(left). ABR thresholds at 4, 8, 12, 16 and 32kHz at P63 of WT, Ndufs4 KO mice administrated with Vehicle or MA-5 (1mg/kg) for 5 weeks(middle). \**p*<0.05, \*\**p*<0.01 vs. WT; #*p*<0.05, ##*p*<0.01 vs. Vehicle. Histological analysis of the mean spiral ganglion cell (SGC) densities in apical, middle, and basal turn of WT and Ndufs4 KO mice after administration of MA-5 (1mg/kg) for 5 weeks, at P62. Data were mean ± SE. \**p*<0.05, \*\**p*<0.01 vs. vehicle; statistics were calculated by one way anova Durnett’s multiple comparison test. **b**, ABR thresholds at 4, 8, 12, 16 and 32kHz at postnatal day 60 (P60) (middle) after administration of vehicle or MA-5 (low dose1.0mg/kg and high dose 10mg/kg) for 3 weeks from P40 when the age-related hearing loss were started \**p*<0.05, \*\**p*<0.01 vs. vehicle, #*p*<0.05, ##*p*<0.01 vs. MA-5 low dose (1.0mg/kg). Histological analysis of SGC densities a in apical, middle, and basal turn of Ndufs4 KO after administration of vehicle control or high dose MA-5 (10 mg/kg) for 3 weeks, at P60. Data were mean ± SE. \**p*<0.05, \*\**p*<0.01 vehicle. Statistical analysis was performed one way anova Durnett’s multiple comparison test. **c**, The auditory brainstem response (ABR) of wild type mice at 4, 8, 12, 16, and 32 kHz at 4 hr and 7 days after the noise exposure in three groups; vehicle (n=6), low dose group (MA-5, 1mg/kg) (n=6), and high dose (MA-5, 10 mg/kg) (n=6). \**p*<0.05, \*\**p*<0.01 vs. vehicle. Histological analysis of survival hair cells in apical, middle, and basal turn of wild type mice after administration of vehicle control or MA-5 (1.0mg/kg) for 14 days, at 7 days after the noise exposure. Data were mean ± SE. \**p*<0.05, \*\**p*<0.01 vs. vehicle; statistics were calculated by one way anova Durnett’s multiple comparison test. **d**, Mouse model of cisplatin ototoxicity. Adult wild mice in three groups; vehicle (n=6), low dose group (MA-5, 1mg/kg) (n=6), and high dose (MA-5, 10 mg/kg) (n=6). Adult wild type mice after administration of vehicle control or MA-5 (10mg/kg) for 9 days, 6 days after 6 cycles of daily cisplatin intraperitoneal injection (4mg/Kg body weight/day) from day4 to day 9 (left).ABR of post cisplatin insult and MA-5 administration (1 and 10 mg/Kg) at 4, 8, 12, 16, and 32 kHz (middle). Data were mean ± SE. \**p*<0.05, \*\**p*<0.01 vs. Vehicle. Histological analysis of survival hair cells, the mean spiral ganglion cell (SGC) densities, thickness of stria vascularis and the mean type IV fibrocyte densities in apical, middle, and basal turn of wild type mice after administration of vehicle control or MA-5 (10mg/kg). Data were mean ± SE.\**p*<0.05, \*\**p*<0.01 vs. vehicle; statistics were calculated by one way anova Durnett’s multiple comparison test. **e**, MA-5 protects against cisplatin-induced ototoxicity in cultured cochlear. Cochlear explants were cultured with various concentration of cisplatin at 0, 5, 7. 5, 10, 15, 20 μM with MA-5 at 10 μM for 48 hours. Viable ocular hair cells (OHCs) counted. \**p*<0.05, \*\**p*<0.01 vs. vehicle (left). Cochlear explants were cultured with cisplatin at 10 μM through various concentrations of MA-5 at 0, 1,5,10, 50 μM for 48hrs. Viable OHCs counted. Data were mean ± SE.\**p*<0.05, \*\**p*<0.01 vs. vehicle (right); statistics were calculated by unpaired two-side t-test.

### MA-5 directly acted on auditory cells, promoting ATP production, and inhibits cell death

Because the blood-labyrinth barrier restricts the entry of most blood-borne compounds into the inner ear, it is essential to confirm whether the drug reaches the cochlea and exerts its effect. We orally administered MA-5 and measured its concentration in the cochlea. The concentration of MA-5 in the cochlea was markedly increased in a dose-dependent manner after oral administration (Fig. 2a). Because MA-5 increases cellular ATP with a reduction in the mitochondrial ROS^10–12^, we collected the organ of Corti and the lateral wall, exposed them to MA-5, and measured the ATP level in the *ex vivo* model (Fig. 2b). MA-5 significantly increased the ATP levels in both the organ of Corti and the lateral wall of the inner ear. Recently, skin fibroblasts from mitochondrial disease patients are useful for measuring mitochondrial function^18^. We previously reported that ATP levels were decreased in patient fibroblasts, and MA-5 amelioraed the reduced levels^12^. Based on the findings that the MA-5 increased ATP levels in the cochlea and improved hearing loss in NDUFS4 KO mice, we applied our findings to humans by collecting fibroblasts from a Leigh syndrome patient with an NDUFS4 mutation and examining the effect of MA-5. MA-5 increased the reduced ATP levels in Leigh syndrome patient cells with NDUFS4 gene mutations in a dose-dependent manner at sub-μM level (Fig. 2c). In addition, in clinical settings, m.1555A>G, mutation is a known risk factor for congenital hearing loss and that increases vulnerability to aminoglycoside^19^. POU4F3 is also one of the most common causes of autosomal dominant progressive hearing loss^20^. Hence, we measured the ATP levels in the fibroblasts from the patients with m.1555A>G or POU4F3 mutations. MA-5 increased the ATP levels in both m.1555A>G and POU4F3 patient skin fibroblasts (Fig. 2c). In addition, fibroblasts from patients with mitochondrial diseases have been reported to be susceptible to cellular reactive oxygen species (ROS) induced by the glutathione synthesis inhibitor L-buthionine-(S,R)-sulfoximine (BSO) ^12,18,21^. This susceptibility provides a useful model for assessing mitochondrial damage and evaluating efficacy of drugs for mitochondrial diseases. In fibroblasts from an NDUFS4 patient, cell death was induced by BSO (Fig. 2d, upper left), and as a result, the LDH level was increased (Fig. 2d, lower left). Under these conditions, the reduced cell survival and elevated LDH levels induced by BSO were reversed by MA-5. Similar to NDUFS4, in m.1555A>G (Fig. 2d, middle upper) and POU4F3 (Fig. 2d, right upper) fibroblasts, cell death was induced by BSO, and the LDH level in m.1555A>G (Fig. 2d, middle lower) and POU4F3 (Fig. 2d, right lower) fibroblasts was increased. Under these conditions, the drug increased cell survival and reduced the elevated LDH levels caused by BSO, suggesting that the drug protects mitochondrial function. These data suggest that MA-5 may be an effective drug for various types of human genetic hearing loss.

**Figure 2.**
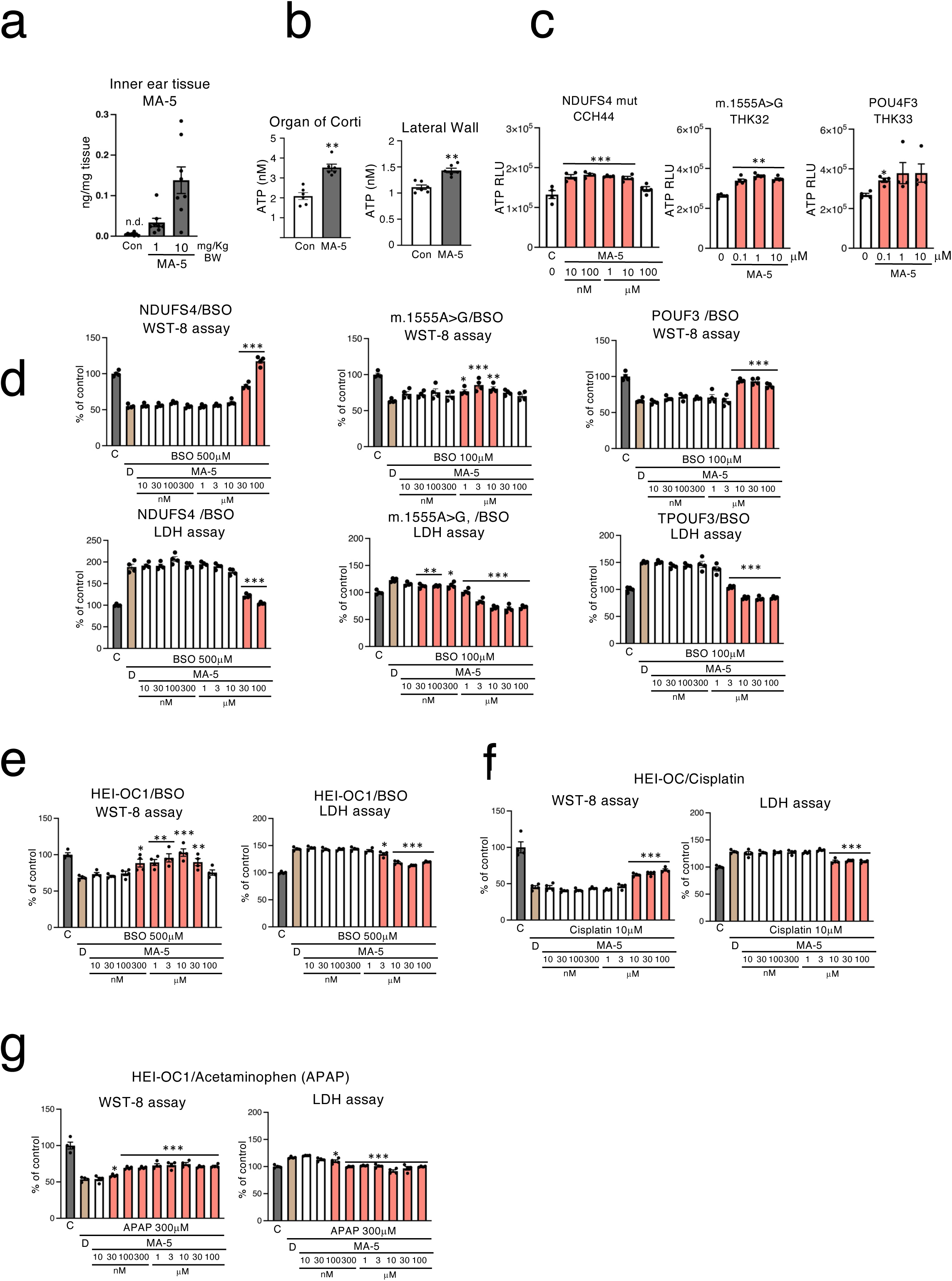
MA-5 enhanced hair cell survival by increasing intracellular ATP levels. **a**, The MA-5 concentration in the cochlea tissue of mice administrated with MA-5. Wild type C57BL6 mice were orally administrated vehicle or MA-5 (10 mg/kg body weight). After 1hr of administration, the cochlea tissues were excised from mice and the concentrations of MA-5 were measured. **b**, The separated parts of cochlear explants were incubated with 10 μM MA-5 for 3hours. ATP contents were measured in tissue homogenate extractions from outer hair cells and lateral walls. Data were mean ± SE. \**p*<0.05, \*\**p*<0.01 vs. control (0.1 % DMSO incubation); statistics were calculated by unpaired two-side t-test. **c**, Intracellular ATP measurement of skin fibroblasts from Leigh syndrome patients (Ndufs4 mutation), m.1555A>G and POU4F3 with various concentration of MA-5 racemate at 10 nM, 100nM, 1 μM,10 μM, 100 μM for 6hr. Data were mean ± SE.\**p*<0.05, \*\**p*<0.01 vs. 0.1 % DMSO control; statistics were calculated by one way anova Durnett’s multiple comparison test. **d,** Cytoprotective effects of MA-5 against cytotoxicity of BSO in skin fibroblasts from Leigh syndrome patients (Ndufs4 mutation), m.1555A>G and POU4F3 patients skin fibroblasts were cultured with BSO at 100μM with various concentrations of MA-5 at 0.1, 0.3, 1, 3, 10, 30n and 100 μM for 48hrs. Cell viability and cytotoxicity were measured by WST-8 assay and LDH assay respectively. Data were mean ± SE.\**p*<0.05, \*\**p*<0.01 vs 0.1% DMSO control; statistics were calculated by one way anova Durnett’s multiple comparison test. **e,** Cytoprotective effects of MA-5 against cytotoxicity of BSO in HEI-OC1 cells. HEI-OC1 cells were cultured with BSO at 500μM through various concentrations of MA-5 at 10, 30, 100, 300nM, 1 μM, 3μM 10 μM and 100 μM for 72hrs. Cell viability and cytotoxicity were measured by WST-8 assay and LDH assay respectively. Data were mean ± SE.\**p*<0.05, \*\**p*<0.01 vs 0.1% DMSO control; statistics were calculated by one way anova Durnett’s multiple comparison test. **f,** Cytoprotective effects of MA-5 against cytotoxicity of cisplatin left in HEI-OC1 cells. HEI-OC1 cells were cultured with cisplatin at 10 μM through various concentrations of MA-5 at 10, 30, 100, 300nM, 1 μM, 3μM, 10 μM and 100 μM for 72hrs. Cell viability and cytotoxicity were measured by WST-8 assay and LDH assay respectively (left). Data were mean ± SE.\**p*<0.05, \*\**p*<0.01 vs 0.1% DMSO control; statistics were calculated by one way anova Durnett’s multiple comparison test. **g,** Cytoprotective effects of MA-5 against cytotoxicity of acetaminophen (APAP) in HEI-OC1 cells. HEI-OC1 cells were cultured with APAP at 500 μM through various concentrations of MA-5 at 10, 30, 100, 300nM, 1 μM, 3μM 10 μM and 100 μM for 48hrs. Cell viability and cytotoxicity were measured by WST-8 assay and LDH assay respectively(right). Data were mean ± SE.\**p*<0.05, \*\**p*<0.01 vs 0.1% DMSO control; statistics were calculated by one way anova Durnett’s multiple comparison test.

To further clarify the effect and mechanism of MA-5 in auditory cells, we used a mouse-derived inner ear cell line, House Ear Institute Organ of Corti 1 (HEI-OC1)^22^. HEI-OC1 is one of the most widely used auditory cell line as a common progenitor for sensory and supporting cells in the organ of Corti^23^. To examine whether HEI-OC1 cells are suitable for drug screening, we firstly performed a sensitivity test to BSO. In HEI-OC1 cells, cell death was induced by BSO accompanied by a corresponding increase in the LDH levels, suitable for drug testing. Under these conditions, MA-5 increased cell survival (Fig. 2e left) and reduced the elevated LDH levels (Fig. 2e right) caused by BSO, suggesting that MA-5 protects mitochondrial dysfunction. We also examined the effects of cisplatin and acetaminophen (APAP) which are known to cause hearing loss. As a result, MA-5 also prevent the cell toxicity induced by cisplatin (survival in Fig. 2f, left and LDH in Fig. 2f, right) and acetaminophen APAP^23^ (survival in Fig. 2g, left and LDH in Fig. 2g, right). These findings indicate that the MA-5 may be effective in preventing or mitigating hearing loss associated with mitochondrial dysfunction and ototoxic agents through enhancement of ATP production and suppression of oxidative stress-induced cell death.

### Differential effects of S-enantiomer and R-enantiomer on the NAD^+^ level

Currently, the phase 1 clinical trial of MA-5 has been finished in Japan (jRCT2031210495) and the phase 2 clinical trial has already been approved by PMDA. MA-5 is a synthetic indole derivative formulated as a racemic mixture. Although clinical trial of MA-5 using the racemic form have been approved, it is generally recommended in drug development that the properties of each enantiomer be evaluated. To clarify the effect of each enantiomer of MA-5, we firstly developed a method for the optical separation and measurement of each MA-5 enantiomer (R-enantiomer and S-enantiomer) using a chiral column and liquid chromatography–tandem mass spectrometry. The chiral column effectively separated both R-enantiomer and S-enantiomer (Fig. 3a, upper), and found that the racemic mixture of MA-5 contain equal amounts of each enantiomer. In some racemic mixtures, one enantiomer can be metabolically converted into the other (chemical inversion/conversion). Therefore, we subsequently administered each enantiomer to rhesus monkeys to investigate whether interconversion occurred between the R- and S-enantiomers *in vivo*. As a result, each enantiomer retained its original configuration in the body of the monkey without chiral interconversion into the other (Fig. 3a, lower panel). This finding suggests that the pharmacological activity of each enantiomer is stereospecific. To elucidate whether the biological effects of MA-5 is due to the R- or S-enantiomer, we initially examined the effect of each enantiomer on the cellular ATP level in HEI-OC1 cells. MA-5 and both enantiomers increased the intracellular ATP level in a dose-dependent manner; however, no difference was observed in potency between MA-5, R-enantiomer, and S-enantiomer (Fig. 3b). To further evaluate the extent of the cytoprotective effect of each enantiomer, fibroblasts derived from a patient with MELAS (mitochondrial encephalomyopathy, lactic acidosis, and stroke-like episodes), one of the more common mitochondrial disorders, carrying the 3243A>G mutation were used.” In 3243A>G patient fibroblasts, BSO- induced cell death was observed as reported^12^ and under the condition, the reduced cell death was significantly ameliorated by each enantiomer, but there was no difference between each enantiomer (Fig. 3c, right; S- enantiomer, left; R-enantiomer).

**Figure 3.**
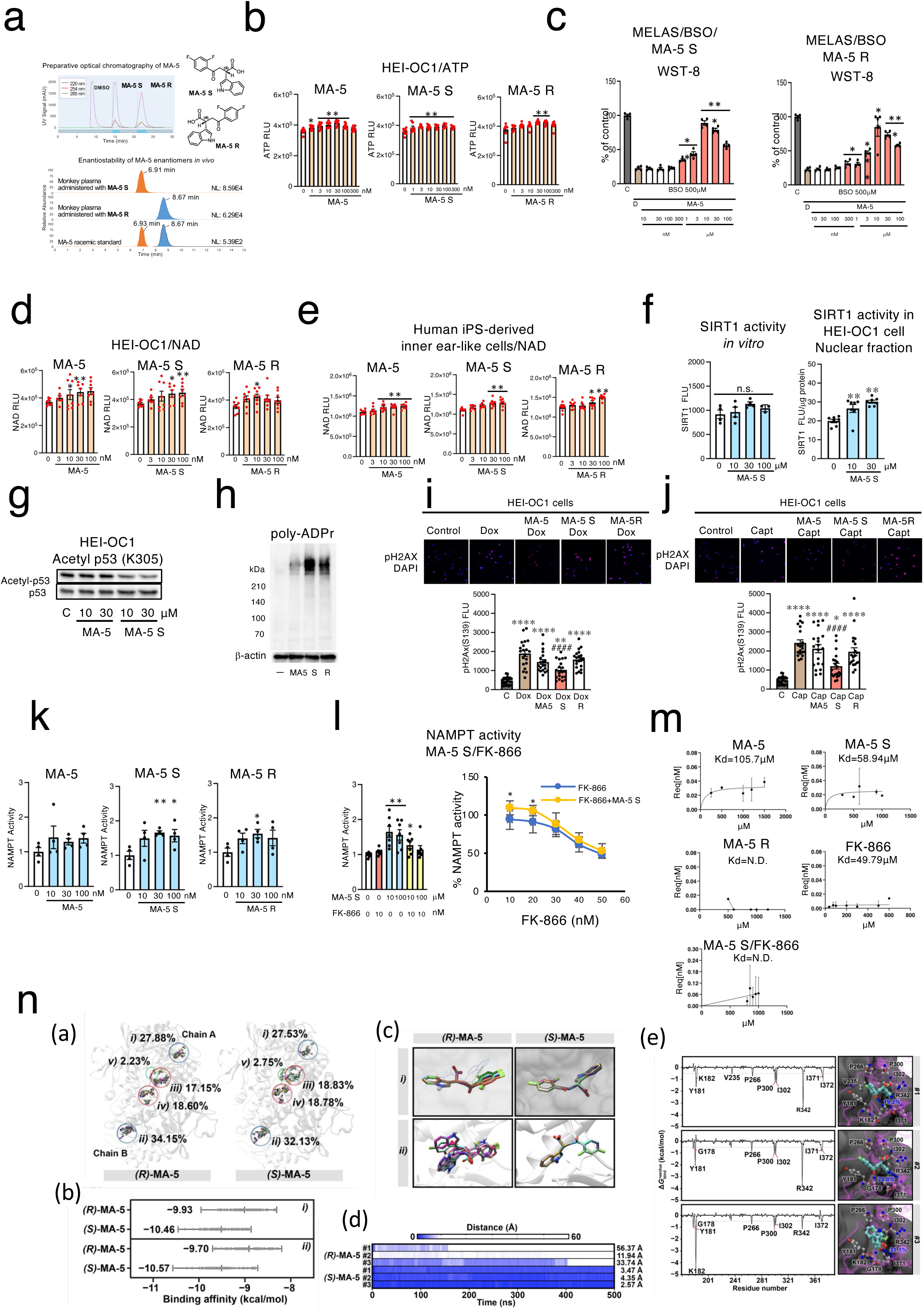
MA-5 acts as a ligand of NAMPT, promoting NAD⁺ accumulation and activating both SIRT and PARP1. **a,** Separation of MA-5 R-enantiomer and S-enantiomer from MA-5 racemic body by the chiral column (upper). The chemical inversion/conversion of R -and S- enantiomer of MA-5 were not observed in a rhesus monkey administered each enantiomer (lower). **b**, Intracellular ATP measurement of HEI-OC1 cells with various concentration of MA-5, S-enantiomer and R-enantiomer at 0, 1, 3,10, 30 and 100nM for 6hr. Data were mean ± SE.\**p*<0.05, \*\**p*<0.01 vs. DMSO control; statistics were calculated by One way anova Durnett’s multiple comparison test. **c**, The cell-protective effect of MA-5 S-enantiomer and R-enantiomer against cytotoxicity of BSO in skin fibroblasts from a MELAS patient. MELAS patient skin fibroblasts were cultured with BSO at 500μM through various concentrations of MA-5 S-enantiomer (left) or R-enantiomer (right) at 10, 30, 100, 300nM, 1 μM, 3μM 10 μM and 100 μM for 72hrs. Cell viability and cytotoxicity were measured by WST-8 assay. Data were mean ± SE.\**p*<0.05, \*\**p*<0.01, \*\*\**p*<0.001 vs 0.1% DMSO control; statistics were calculated by One way anova Durnett’s multiple comparison test. **d,** Intracellular NAD ^+^ measurement of HEI-OC1 cells. HEI-OC1 cells were treated with 0.1 % DMSO or MA-5 racemate, MA-5 S-enantiomer and R- enantiomer at 3,10, 30 and 100nM for 6hr. Data were mean ± SE. \**p*<0.05, \*\**p*<0.01 vs. DMSO control; statistics were calculated by One way anova Durnett’s multiple comparison test. **e**, Intracellular NAD ^+^ measurement of human iPS-induced inner ear cells. Human iPS-induced inner ear cells were treated with 0.1 % DMSO or MA-5 racemate, MA-5 S-enantiomer and R-enantiomer at 3,10, 30 and 100nM for 6hr. Data were mean ± SE. \**p*<0.05, \*\**p*<0.01 vs. DMSO control; statistics were calculated by One way anova Durnett’s multiple comparison test. **f,** The effects of S-enantiomer on SIRT1 enzymatic activity. The effects of S- enantiomer on SIRT1 deacetylase activity *in vitro* at 10,30 and 100 μM (left). The effects of S-enantiomer on SIRT1 deacetylase activity in cell nuclear fraction extract from HEI-OC1 cells treated with S-enantiomer at 10 and 30 μM for 3hr (right). Data were mean ± SE.\**p*<0.05, \*\**p*<0.01 vs. DMSO control; statistics were calculated by One way anova Durnett’s multiple comparison test. **g,** The acetylation of p53 (K305) was reduced by MA-5 S enantiomer. The effects of S-enantiomer on the acetylation of p53 (K305) in whole cell extraction from HEI-OC1 cells treated with MA-5 racemate and S-enantiomer at 10 and 30 μM for 24hr. **h**, The cellular localization of PARylation in HEI-OC1 cells treated with 0.1 % DMSO or MA-5 at 10μM for 24hr. (left). Poly-ADP ribosylation of PARP protein in HEI-OC1 cells treated with 0.1 % DMSO, MA-5 racemate, S-enantiomer and R-enantiomer at 30 100μM for 24hr. **i**, Immunocyto-staining of H2AX phosphorylated (γH2AX) of HEI-OC1 cells treated with 0.1 % DMSO, doxiorubicin (Dox) at 0.3μM, MA-5, S-enantiomer and R-enantiomer at 30 μM for 24hr. Representative (Upper). Florescence intensities of γH2AX immunocytostaining were measured (Lowere). Data were mean ± SE. *p*<0.05, ***: *p*<0.0001 vs. control. # *p*<0.05, ####*p*<0.0001 vs. Dox 0.3μM; statistics were calculated by One way anova Durnett’s multiple comparison test. **j**, Immunocytostaining of H2AX phosphorylated (γH2AX) of HEI-OC1 cells treated with 0.1 % DMSO, camptothecin (Capt) at 1μM, MA-5, S-enantiomer and R-enantiomer at 30 μM for 24hr. Representative (Upper). Florescence intensities of γH2AX immunocytostaining were measured (Lowere). Data were mean ± SE. *p*<0.05, ***: *p*<0.0001 vs. control. # *p*<0.05, ####*p*<0.0001 vs. Capt 1μM; statistics were calculated by One way anova Durnett’s multiple comparison test. **k**, The NAMPT enzymatic activity *in vitro* treated with MA-5, S-enantiomer and R- enantiomer at 3,10, 30 and 100nM for 1hr. Data were mean ± SE. \**p*<0.05, \*\**p*<0.01 vs. DMSO control; statistics were calculated by One way anova Durnett’s multiple comparison test. **l,** The NAMPT enzymatic activity *in vitro* treated with S-enantiomer at 10 and 100μM with or without selective NAMPT inhibitor FK-866 at 10nM (**left**). Data were mean ± SE.\**p*<0.05, \*\**p*<0.01 vs. DMSO control; statistics were calculated by One way anova Durnett’s multiple comparison test. The Inhibitory curve on NAMPT activity with various concentration of FK-866 at at 10, 20, 30, 40 and 50nM (**right**). The Inhibitory curve on NAMPT activity with FK-866 was shifted by addition of MA-5 S-enantiomer at 100 μM (**right**). \**p*<0.05, \*\**p*<0.01 vs. FK-866 alone inhibitory curve; s statistics were calculated by unpaired two-side t-test. **m**, The binding between NAMNPT and MA-5, MA-5 S-enantiomer and R-enantiomer and FK-866 were quantitatively analyzed using Biolayer Interferometry (BLI). **n**, In silico study of Mitochonic acid 5 binding to NAMPT. (a) Possible binding sites on NAMPT of MA-5 S-enantiomer and R-enantiomer. (b) The binding interaction scores of MA-5 S-enantiomer and R-enantiomer at sites (*i*) and (*ii*). (c) The binding conformations of MA-5 S-enantiomer and R-enantiomer at sites (*i*) and (*ii*). (d) the conformational dynamics of ligand binding of MA-5 S-enantiomer and R-enantiomer. (e)

### S-enantiomer increased cellular NAD^+^ level

Our previous studies have highlighted the importance of not only ATP but also nicotinamide adenine dinucleotide (NAD^+)^, and sirtuins (SIRTs) are important in maintaining the hearing function^6^. The reduction of NAD^+^ is associated with the progression of ARHL^24^, cisplatin-mediated hearing damage^25^ and NIHL^26^, suggesting that increasing the NAD^+^ levels protects hearing capacity. Therefore, we next examined the effects of MA-5 and each enantiomer on NAD^+^ levels. When the effect of MA-5 on NAD⁺ levels was evaluated, the racemic mixture induced a slight increase in cellular NAD⁺. In contrast, the S-enantiomer significantly elevated NAD⁺ levels in a dose-dependent manner (Fig. 3d), whereas the R-enantiomer had no significant effect. These results suggest that the beneficial effect of the S-enantiomer on NAD⁺ production may be attenuated by the presence of the R- enantiomer, leading to only a modest increase in NAD⁺ levels when the racemic mixture MA-5 is administered. Recently, we established human iPS cell-derived inner ear cells, providing a novel platform for evaluating therapeutic strategies for progressive hearing loss. So, we next examined the effects of MA-5 and its enantiomers on the NAD^+^ levels in human iPS-derived inner cells as reported^27^. MA-5 and the S-enantiomer increased the NAD^+^ levels in human iPS-derived inner cells, while the R-enantiomer only increased the NAD^+^ levels at higher doses (Fig. 3e). These data further suggest that S-enantiomer not only increases cellular ATP levels but also enhances NAD⁺ levels. It is reported that the NAD^+^ concentration regulates the activity of its consuming enzymes, SIRT^28^ and poly(ADP-ribose) polymerase 1 (PARP1)^29^. Considering that the S-enantiomer increased the NAD^+^ levels, we next evaluated the effects of MA-5 and enantiomers on SIRT1 and PARP1 enzyme activities. Without cellular extraction, neither MA-5 nor the enantiomers affected the isolated SIRT1-3 enzymatic activity (Fig. 3f, left, Extended Data Fig. 2a). However, when S-enantiomer was exposed to HEI-OC1 cells, the SIRT1 activity of nuclear fraction-cell lysate significantly increased (Fig. 3f, right), These data suggest that the S-enantiomer mobilized the intracellular SIRT1 activity without direct enzyme activation. Because SIRT1 deacetylates P53 in a NAD^+^-dependent manner and deacetylation of p53 prevents the cellular senescence and apoptosis caused by DNA damage and stress^30^, we next examined the acetylation level of P53. Consequently, acetylation of P53 (K305) was decreased by S-enantiomer compared with MA-5 (Fig. 3g). These data indicate that the S-enantiomer not only increased the NAD^+^ levels but also increased the SIRT1 enzymatic activity and may exerts cytoprotective and DNA repair effects through the deacetylation of p53.

### S-enantiomer modulated the PARP activity, following the reduction of DNA damage

In the context of DNA damage repair, poly [ADP-ribose] polymerase 1 (PARP-1) functions as a DNA damage sensor, and NAD⁺ serves as a co-substrate for PARPs^31^. Hence, we subsequently examined the effects of MA-5 and enantiomers on poly(ADP-ribosyl)ation (PARylation), and the PARP1 activity. The S-enantiomer induced significantly higher levels of DNA PARylation compared to the MA-5 and the R-enantiomer (Fig. 3h). On the other hand, no change was observed in the Su of lysine (K) or the acetylation of lysine (Extended Data Supplementary Fig. 2b). Because S-enantiomer did not have any direct effect on the PARP1 enzyme activity (Extended Data Fig. 2c), these data further suggest that PARylation induced by the S-enantiomer is a consequence of the increased NAD^+^ levels rather than a direct enzymatic activation of PARP1. Because DNA damage activates PARP1^32^, we next examined whether PARP1 activation by S-enantiomer is a result of DNA damage. In HEI-OC1 cells, the DNA-damaging agent doxorubicin (DOX) induced DNA damage and increased the phosphorylation of γH2AX, a sensitive markers of DNA damage. However, under the condition, neither MA-5 nor enantiomers resulted in an increase in γH2AX, but S-enantiomer significantly decrease DOX-induced γH2AX (Fig. 3i). Similarly, the expression level of the γH2AX expression induced by the DNA-damaging agent camptothecin (CPT) in HEI-OC1 cells was also mitigated by S-enantiomer (Fig. 3j). These data suggest that the S-enantiomer increased PARP1 activity without damaging DNA and, moreover, the S-enantiomer showed a DNA-protective effect through SIRT and PAPR1 activation.

### S-enantiomer increased NAD^+^ as an NAMPT activator

NAD^+^ is synthesized from nicotinamide (NAM) through a two-step enzymatic reaction; NAM is converted into nicotinamide mononucleotide (NMN) primarily by nicotinamide phosphoribosyltransferase (NAMPT), and nicotinamide mononucleotide adenylyl transferases 1 and 3 (NMNAT1 in the nucleus and NMNAT3 in the mitochondria) catalyze the subsequent conversion of NMN to NAD⁺. Among these steps, nicotinamide phosphoribosyltransferase (NAMPT) serves as the rate-limiting enzyme in the NAD^+^ salvage pathway and is essential for NAD^+^ biosynthesis in most mammals^31^. So, we next examined the effect of MA-5 and enantiomers on NAMPT activity. Racemic MA-5 has less effect on NAMPT activity. On the other hand, S-enantiomer caused the activation of human NAMPT-mediated NMN production at the nM level (Fig. 3k). A weak effect on the NAMP activity was also seen by R-enantiomer. In addition, this NAMPT-activating ability of S-enantiomer was abolished by a NAMPT inhibitor, FK-866^33^ (Fig. 3l, left), in a dose-response manner (Fig. 3l, right). On the other hand, NAM mononucleotide adenylyl transferase 1 (NMNAT1) and NMNAT3 synthesize NAD**^+^** from nicotinamide mononucleotide (NMN). Although we examined the effects of MA-5 and its enantiomers on NMNAT1 and NMNAT3 enzyme activities (Extended Data Fig. 3), neither MA-5 nor its enantiomers affected the activities of NMNAT 1 or NMNAT3. These results suggest that the S-enantiomer may act as a NAMPT activator to increase NAD^+^. To further examine the interaction between NAMNPT and the S-enantiomer, the binding capacity of NAMPT was analyzed using Biolayer Interferometry (BLI). Recombinant human NAMPT were purified (Extended Data Fig. 4) and mobilized on avidin tips, and the sensors were exposed to ligand solutions and investigated the binding of MA-5 and its enantiomers to NAMPT (Fig. 3m). The binding curves obtained with increasing concentrations of MA-5 showed binding, with an estimated *Kd* of 105.7 μM. Moreover, the S-enantiomer exhibited more specific binding to NAMPT, with a *Kd* of 58.94 μM. Conversely, no binding affinity was observed for the R-enantiomer. As the NAMPT activation of the S-enantiomer was inhibited by FK866 (Fig. 3l), we further investigated whether FK866 could inhibit the S-enantiomer binding to NAMPT.

FK866 showed significant binding to NAMPT, with an estimated *Kd* of 49.79 μM, and its presence completely inhibited the S-enantiomer’s binding to the recombinant NAMPT protein. These results suggest that the S-enantiomer shares an allosteric FK866 binding site with NAMPT. To further predict the potential binding sites of NAMPT and its preferred stereoisomer, we employed the blind docking with the k-means clustering (BDK) technique ^34^. Both enantiomers were docked to the entire protein structure over 100 runs using AutoDock Vina 1.2.3. A total of 4,000 ligand conformations revealed five possible binding sites on NAMPT with varying probabilities (Fig. 3n (a)). This indicated that sites (*i*) and (*ii*), corresponding to the active sites of NAMPT, were the most likely binding sites for the S-enantiomer. Consequently, focused docking was performed at these sites to refine the previous docking results. The lowest binding interaction scores at sites (*i*) and (*ii*) were −9.70 to −9.93 kcal/mol for R-enantiomer and −10.46 to −10.57 kcal/mol for S-enantiomer (Fig. 3n (b)). Notably, the binding conformation of the S-enantiomer showed consistency at sites (*i*) and (*ii*), suggesting that this conformation is preferred for the ligand-binding mechanism (Fig. 3n (c)). To explore the conformational dynamics of ligand binding, both S- and R-enantiomer complexes bound to the active site of NAMPT were subjected to 500-ns MD simulations with three replications. In the R enantiomer system, ligand dissociation from the protein occurred at approximately 154 and 411 ns in system #1 and system #3, respectively (Fig. 3n (d)). In system #2, the ligand rapidly dissociated from the binding pocket and relocated to the protein surface. Based on these observations, only the S-enantiomer system was selected for further analysis of the binding strength and interactions using the molecular mechanics-generalized born and the surface area solvation method. The S-enantiomer demonstrated stable binding to the active site of NAMPT with binding energies of −25.76±0.53 kcal/mol (#1), −23.10±0.36 kcal/mol (#2), and −21.93±0.46 kcal/mol (#3). Among the three replicates, the ligand conformations in #1 and #2 were almost identical. The stabilization of the S-enantiomer involved π-π stacking between Y181 and the 4-(2,4-difluorophenyl) ring, along with hydrogen bonding interactions between the nitrogen of the indole ring and K182 (∼19.2%) in #1 and R342 (∼23.8%) in #2. In #3, a distinct binding pattern was observed, where the 4-(2,4-difluorophenyl) ring of the ligand folded back toward its indole ring in a globular-like structure, forming hydrogen bonds with K182 (∼31.1%) and plugging into the active site (Fig. 3n (e)). These computational studies suggest that the S-enantiomer binds stably to NAMPT, unlike the R-enantiomer. This preference is supported by the favorable binding energies and interactions with key residues. Variations in ligand conformation were observed, emphasizing the importance of stereoisomers in ligand-binding study and highlighting the therapeutic potential of the S-enantiomer in NAMPT-targeted interventions. These data suggest that the S-enantiomer acts as an agonist of NAMPT and enhances NAD^+^ following the activation of the SIRT1 and PAPR activities.

### S-enantiomer increased SIRT1-7 proteins

The mammalian family of SIRTs consists of seven enzymes, SIRT1-7 and SIRT activities could be critically influenced by physiological changes in the NAD^+^ levels^24^. However, to date, there is no evidence to support that increased NAD⁺ levels directly upregulate the expression of sirtuin proteins. So, we next investigated the changes in SIRT1-7 expression levels by MA-5 and its enantiomers. Quantitative PCR revealed that neither MA-5, S- enantiomer, nor R-enantiomer affected the SIRT1-7 mRNA levels (Fig. 4a). In contrast, surprisingly, the protein expression levels of SIRT1-7 were significantly increased by the S-enantiomer in a dose-dependent manner (Fig. 4b). On the other hand, no significant increase in the SIRT1-7 protein was detected with R-enantiomer. Consequently, a moderate upregulation of SIRT1-7 proteins was induced by MA-5 (Fig. 4b). In human iPS- derived inner ear cells, the S-enantiomer also increased the SIRT1-SIRT7 protein expression levels (Fig. 4c). To confirm this *in vivo*, we examined the effects in mouse cochlea. Because we do not have enough amount of S- enantiomer for *in vivo* experiment, we administer MA-5 (10 μg/Kg) to normal mouse and the expression levels of SIRT1 and SIRT3 was examined. The expression level of SIRT1 protein was increased by MA-5 in mouse cochlea (Fig. 4d, upper). Histological analysis using spinning-disk confocal technology^35^ also showed the increased SIRT1 expression in the stria vascularis of the cochlea (Fig. 4d, lower). These findings confirm that the S-enantiomer not only increased NAD^+^ production but also increased the protein expression levels of SIRT1-7 *in vitro* and *in vivo* without changing the transcriptional regulation. The differential expression of SIRT1 and the absence of detectable SIRT3 in the cochlea may be attributable to the administration of the racemic form of MA-5.

**Figure 4.**
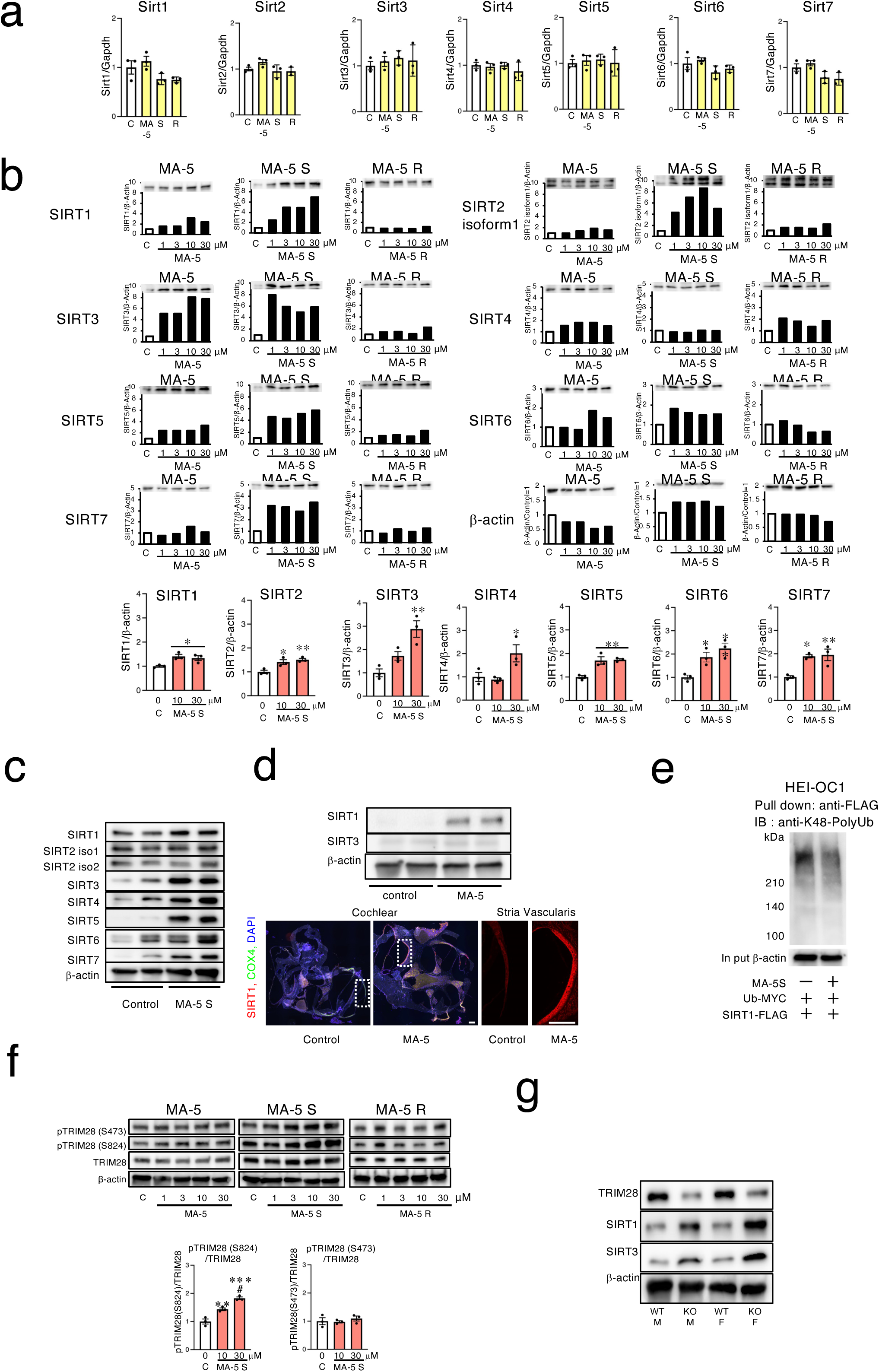
MA-5 upregulates SIRT proteins via a TRIM28-dependent mechanism. **a,** The mRNA levels of Sirt1-Sirt7 in HEI-OC1 cells treated with MA-5, S-enantiomer and R-enantiomer at 30 μM for 24hr. The quantitative real time PCR of mouse Sirt1-Sirt7 were performed and standardized by internal standard of mouse Gapdh. Data were mean ± SE.; statistics were calculated by One way anova Durnett’s multiple comparison test. **b,** The protein expression levels of SIRT1-SIRT7 in HEI-OC1 cells treated with MA-5, S-enantiomer and R-enantiomer at 1, 3, 10, 30 μM for 24hr. Protein expression levels of SIRT1-7 were standardized by protein expression level of β-actin. \**p*<0.05, \*\**p*<0.01 vs. control; statistics were calculated by One way anova Durnett’s multiple comparison test. **c**, The protein expression level of all SIRT1-SIRT7 in human iPS-human inner ear cell-like-cells treated with MA-5 S-enantiomer at 30 μM for 24hr. **d**, The expression level of SIRT1 protein in mouse cochlear with or without MA-5 treatment (10mg/Kg). Large scan images of cochlea and typical images of stria vascularis. Scale bar, 100 µm. **e**, K48-linked polyubiquitin of SIRT1 protein. The protein of SIRT1 with FLAG-tag and Ubiquitin with Myc tag were over expressed in HEI-OC1cells by expression vector transfection. HEI-OC1 cells were treated with MA-5 S-enantiomer at 0 and 30 μM for another 24hr after 48hr from expression vector transfection. Myc-tag beads pull down protein extractions were accessed by Western blot analysis with anti-K48-linked polyubiquitin specific anti body. **f**, The phosphorylation of TRIM28(Ser 824) and Phosphorylation of TRIM28(Ser 473) with TRIM28 hole protein expression in HEI-OC1 cells treated with MA-5 racemate, MA-5 S-enantiomer and R-enantiomer at 1, 3, 10, 30 μM for 24hr. Lower panel, the quantitative analysis of TRIM28(Ser 473) and TRIM28(Ser 824). \**p*<0.05, \*\**p*<0.01 vs. control; statistics were calculated by One way anova Turkey’s multiple comparison test. **g**, The SIRT1, SRIT3 and TRIM28 protein expression skin fibroblasts cells form two TRIM28-hetero knockout mice (male and female) and their littermate two wild type mice (male and female).

### Post-translational modification by TRIM28 regulating SIRT1-7 protein stability

As MA-5 did not affect SIRT1-7 expression at the transcriptional level, we next examined its degradation pathway. Recently, Huang et al. reported that SIRT3 protein expression was enhanced by MA-5 mediated by the adenosine monophosphate-activated protein kinase (AMPK) pathway^36^. To determine the involvement of the AMPK pathway in the S-enantiomer-induced enhancement of SIRT1-7 protein expression, we used the AMPK pathway inhibitor compound C. Compound C had no effect on the S-enantiomer-induced expression of SIRT1, 2, and SIRT 4– SIRT7 proteins except SIRT3 (Extended Data Fig. 5). These findings suggested that the SIRT protein upregulation mechanism of S-enantiomer was not mainly dependent on the AMPK pathway. It is reported that the primary mechanism regulating the expression of SIRT proteins is the ubiquitination and K48-linked polyubiquitination promotes their degradation^37^. Therefore, we examined the effects of the S- enantiomer on SIRT1 ubiquitination. Pulldown analysis of ubiquitinated SIRT1 revealed that the SIRT1 ubiquitination level was decreased by S-enantiomer in HEI-OC1 cells (Fig. 4e). On the other hand, the other E3 ubiquitin ligase inhibitors thalidomide, lenalidomide, and NSC 66811 did not affect the SIRT1 protein expression (Extended Data Fig. 6). These data suggest that protein degradation of SIRT1 is modulated by S-enantiomer. Tripartite motif-containing 28 (TRIM28, also known as Krüppel-associated box-associated protein 1 (KRAB1) or transcriptional intermediary factor 1β (TIF1β) contains several phosphorylation sites, among which serine 473 (S473) is associated with stress and inflammatory responses, whereas serine 824 (S824) is phosphorylated in response to DNA damage and facilitates chromatin relaxation, thereby promoting DNA repair^38^. Recently, Ouyang et al. reported that phosphorylation S824 of TRIM28 decreases its ubiquitination activity and increases the SIRT1 protein expression^39^. So, we examined the phosphorylation status of TRIM28 in response to the MA-5 and its enantiomers in HEI-OC1 cells. S-enantiomer promoted TRIM28 phosphorylation of Ser824 but not Ser473, in a dose-dependent manner (Fig. 4f). In contrast, the R-enantiomer had no effect on the TRIM28 phosphorylation. To further clarify the relationship between TRIM28 and SIRT proteins, we generated TRIM28 knockout (KO) mice^40^ and examined the expression levels of TRIM28, SIRT1, and SIRT3 in the skin fibroblasts (Fig. 4g). Since homozygote is lethal, we used heterozygous KO mice fibroblast for the experiment. As a result, the protein level of TRIM28 was decreased in KO fibroblasts and vice versa, the SIRT1 and SIRT3 protein levels were increased in TRIM28 KO fibroblasts. These results suggest that the S-enantiomer phosphorylates Ser824 of TRIM28 and upregulates SIRT1-7 protein expression by reducing its ubiquitination. Generally, TRIM28 phosphorylation occurs in response to DNA damagel^41^, therefore, it is suspected that the phosphorylation of TRIM28 by the S-enantiomer may occur by DNA damage. However, here, already showed that S-enantiomer induces PRPR1 activity and SIRT1 activation and has DNA protective effects (Fig. 3i and j). In addition, during the clinical development process of MA-5, several tests for determining the genotoxic tests including the Ames test, micronucleus assay at clinical trial good manufacturing practice (GMP) level has been performed and confirmed that MA-5 is not DNA mutagenic. These data indicate that activation of phosphorylation Ser824 of TRIM28 by the S-enantiomer is not due to the DNA damage.

### Non-DNA-damaging DNA-PK activation by S-enantiomer

Ataxia Telangiectasia Mutated (ATM) phosphorylates TRIM28 at serine 824 (S824), leading to the dissociation of TRIM28 from heterochromatin and subsequent chromatin decondensation. This relaxation of chromatin structure facilitates the recruitment of DNA repair proteins to sites of damage, thereby promoting efficient DNA repair^42^. DNA damage induces rapid autophosphorylation of Ser 1981 of ATM that causes dimer dissociation and initiates cellular ATM kinase activity^43^. ATM also phosphorylates itself at S1981 and promotes dimer-to-monomer transition from an inactive dimer to an active monomer in response to DNA damage^44^. To determine whether TRIM28 phosphorylation by S-enantiomer was dependent on ATM activation, we investigated the phosphorylation status of ATM (S1987). However, S-enantiomer did not affect the phosphorylation of ATM (Fig. 5a). We next investigated whether KU-60019, a specific inhibitor of ATM/ATR, influences the effect of S-enantiomer. As a result, the upregulation of SIRT1/3 protein expression by S-enantiomer was not suppressed by KU-60019 (Fig. 5b). To further confirm that the phosphorylation of TRIM28, we used ATM knockdown HEI-OC1 cells. In ATM knockdown HEIOC-1 cells, S-enantiomer still increased the phosphorylation of TRIM28 (S824) (Fig. 5c). These data suggest that S-enantiomer activates TRIM28 phosphorylation different from DNA damaging signal.

**Figure 5.**
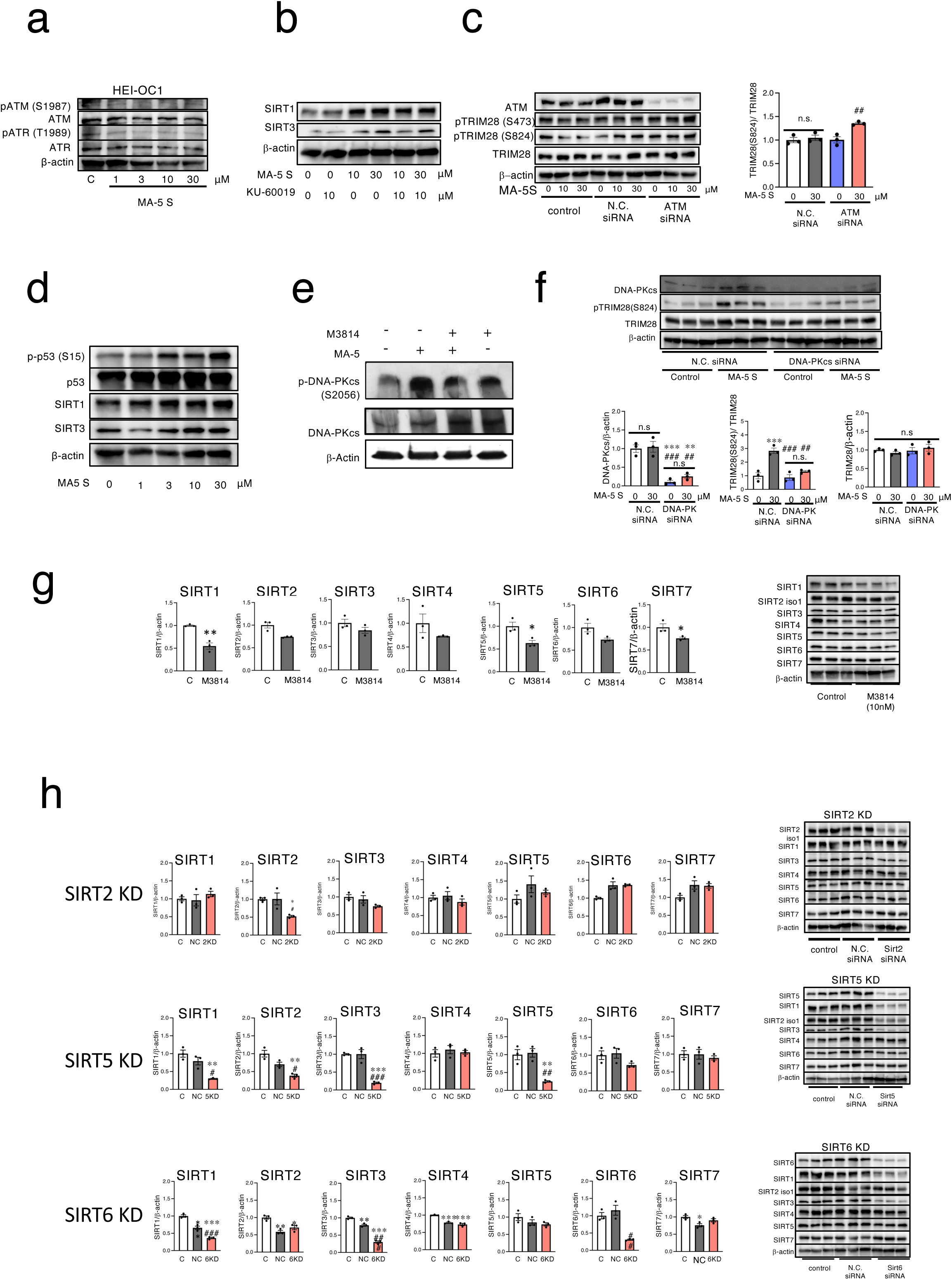
MA-5 modulates DNA-PK phosphorylation without inducing DNA damage. **a**, The phosphorylation of ATM(Ser 1987), ATM whole protein in HEI-OC1 cells treated with MA-5 S- enantiomer at 0, 3,10, 30 μM for 24hr. **b**, The SIRT1 and SRIT3 protein expression in HEI-OC1 cells treated with S-enantiomer at 0 and 30 μM with or without ATM inhibitor KU-60019 at 10μM for 24hr. **c**, Phosphorylation of TRIM28(Ser 824), Phosphorylation of TRIM28(Ser 473) TRIM28 hole protein and ATM protein expression in in HEI-OC1 cells after ATM siRNA knock down. Right panel, the quantitative analysis of TRIM28(Ser 824) phosphorylation. \**p*<0.05, \*\**p*<0.01 vs. control; statistics were calculated byOne way anova Turkey’s multiple comparison test. **d**, Phosphorylation of p53(Ser 15) expression and p53 whole protein in HEI-OC1 cells treated with MA-5 racemate at 0, 1, 3,10, 30 μM for 24hr. **e**, The phosphorylation of DNA-PKcs(Ser 2056) and DNA-PKcs whole in HEI-OC1 cells treated with S-enantiomer with or without DNA-PK inhibitor, M3814 (10nM). **f**, The phosphorylation of TRIM28(Ser 824) and TRIM28 hole protein and DNA-PKcs hole protein expression in in HEI-OC1 cells after DNA-PKcs siRNA knock down procedure. Control (Lipofectamine-RNAiMAX regent alone without siRNA), Negative-Control (NC) siRNA transfected HEI-OC1cells were incubated for 48hr and treated with MA-5 S enantiomer at 30 μM for another 24hr. Data were mean ± SE. \**p*<0.05, \*\**p*<0.01 vs. DMSO control; statistics were calculated by One way anova Turkey’s multiple comparison. **g**, DNA-PK inhibitor reduced the S-enantiomer-induced SIRT1 expression. The protein expression level of all SIRT1-SIRT7 in HEI-OC1 cells treated with 0.1 % DMSO or DNA-PK inhibitor M3814 at 10nM for 24hr. Data were mean ± SE. \**p*<0.05, \*\**p*<0.01 vs. DMSO control; statistics were calculated by unpaired two-side t-test. **h**, The he protein expression of SIRT1-7 when SIR2, SIRT5, and Sirt6 were knocked down. Data were mean ± SE. \**p*<0.05, \*\**p*<0.01 vs. DMSO control; statistics were calculated by one way anova Turkey’s multiple comparison.

DNA-dependent protein kinase DNA-PK is a critical component in the repair of DNA double-stranded breaks and DNA-PK is also essential for TRIM28(Ser824) phosphorylation under stress conditions^45^. In addition, It is well known that DNA-PK is responsible for inducing p53(Ser15) phosphorylation^46^. To further clarify the mechanism of Ser824 phosphorylation of TRIM28, the phosphorylation level of P53(Ser15) was examined. As a result, S-enantiomer enhanced the phosphorylation of P53(Ser15) in HEI-OC1 cells in a dose response manner (Fig. 5d), suggesting the S-enantiomer activated of DNA-PK. In addition, it is well known that phosphorylation of DNA-PKcs at the S2056 ensures it activity and is needed for efficient DNA-DSB repair^47^. Therefore, we next investigated whether the S-enantiomer affects DNA-PK phosphorylation. As a result, S-enantiomer enhanced the phosphorylation of the S2056 of DNA-PK, and the phosphorylation of S2056 was inhibited by DNA-PK inhibitor M3814 (Nedisertib)^48^ (Fig. 5e). To further confirm this, we downregulated DNA-PK in HEIOC-1 cells using siRNA and examined the effect of the S-enantiomer on TRIM28 phosphorylation. As a result, the phosphorylation of TRIM28 S824 induced by S-enantiomer was canceled in DNA-PK knockdown HEIOC-1 cells (Figure 5f). To further clarify the role of DNA-PK on S-enantiomer-induced SIRT1-7 upregulation, we examined the effect of M3814 on SIRT1-SIRT7 protein expression level. As shown in Figure 5g, low dose of M3814 (10 nM) inhibited the SIRT1, SIRT5 and SIRT7 protein expressions. These data suggest that the S-enantiomer accelerates DNA-PK phosphorylation and activates DNA-PK, following phosphorylates TRIM28(S824) and maintains SIRTs protein expression. Docking calculations using MOE were performed to determine the binding mode of S-enantiomer to DNA-PKcs. The docking of S-enantiomer was carried out against the same binding pocket as M3814. We utilized a known structure (PDB ID: 7OTY) as a complex of DNA-PK and M3814. The binding mode of S-enantiomer was observed to form hydrophobic and electrostatic interactions (Extended Data Fig. 7a, left). No steric hindrance was observed between the docked S-enantiomer and the ligand binding pocket of DNA-PKcs (Extended Data Fig. 7a, right). The interaction manner of the S-enantiomer to DNA-PK was likely similar to that of M3814, with the relatively hydrophobic moiety of the compound located deep in the protein pocket ((Extended Data Fig. 7b). These data further suggest that S-enantiomer is a partial ligand of DNA-PKcs.

Furthermore, to date, only limited evidence exists to support that one sirtuin directly interacts with and regulates the expression of another. For instance, administration of the SIRT1 activator SRT1720 has been shown to increase SIRT3 expression and promote mitochondrial biogenesis^49^. Sirt6 and Sirt3, regulate each other’s activity and protect the heart from developing obesity-mediated diabetic cardiomyopathy^50^. In addition, in the brains of SIRT6-deficient mice, a significant reduction in the expression levels of SIRT3 and SIRT4 has been reported, accompanied by impaired mitochondrial function^51^. These findings suggest that there may be some level of interaction among sirtuins. Because S-enantiomer increased SIRT1-7 proteins expression, we further assumed that up-regulation of SIRTs protein synergistically increased SIRTs proteins. To clarify this sirtuin-sirtuin interaction, we examined each sirtuin knockdown experiment (Fig. 5h). SIRT5 knockdown led to a decrease in the protein expression levels of SIRT1, SIRT2, and SIRT3, as well as SIRT5 itself without changing SIRT1-3 mRNA expression. SIRT6 knockdown led to a reduction in the protein expression levels of SIRT1 and SIRT3. Concerning SIRT2 knockdown, no change was shown in the SIRT1-7 protein level, suggesting that sirtuin-sirtuin interaction. These data suggested that interactions among sirtuin proteins may play a role in regulating each other’s expression, and that the upregulation of certain sirtuin(s) by MA-5 may also contribute to the increased expression of other members of the sirtuin family. Overall, S-enantiomer modulates DNA-PK activity, following TRM28 phosphorylation, and increased the protein level of SIRT1-7. S-enantiomer may also increase SIRTs protein expression by modulating sirtuin-sirtuin interaction.

### MA-5 mimics the pluripotency signatures

To investigate the effects of MA-5 on cochlear transcriptomes, we conducted an RNA sequencing (RNA-seq) analysis of cochlear samples from WT and Ndufs4 KO mice. MA-5 treatment of NDUFS4 mice resulted in the identification of 621 DEGs, of which 422 were upregulated and 199 were downregulated (Fig. 6a). The DEGs were visualized using a heat map and in Ndufs4 KO mice, 430 DEGs were identified, with 242 genes upregulated and 188 downregulated following MA-5 treatment (Fig. 6a. left). A pathway analysis of the DEGs identified from the RNA-seq data revealed that in Ndufs4 KO mice, MA-5 administration led to the downregulation of pathways linked to the epithelial mesenchymal transition (EMT), estrogen response, myogenesis, tumor necrosis factor (TNF)-α signaling via nuclear factor κB (NF-κB), apical junction, hypoxia, KRAS signaling, coagulation and ultraviolet (UV) response (Fig. 6c). On the other hand, pathways enriched for genes upregulated by MA-5 in NDUFS-4 KO mice cochlear included G2-M checkpoint, E2F targets, heme metabolism, mitotic spindle, Myc targets V1, xenobiotic metabolism, interferon γ response, DNA repair and late estrogen response (Fig. 6b). In HEI-OC1 cells, 131 DEGs were identified, with 56 genes upregulated and 75 genes downregulated by MA-5 treatment (Fig. 6c. left). The DEGs were visualized using a heat map (Fig. 6c, right). A pathway analysis of the DEGs identified from the RNA-seq data revealed that in HEI-OC1 cells, similar to NDUFS4 KO mice cochlear, EMT, mitotic spindle, coagulation, cholesterol homeostasis, KRAS signaling, myogenesis, TNF-α signaling NF-κB, TGF-β signaling, UV response down, and apoptosis were decreased (Fig. 6d). On the other hand, bile acid metabolism, adipogenesis, xenobiotic metabolism, heme metabolism, oxidative phosphorylation, UV response, Myc targets (V1), glycolysis, p53 pathway and fatty acid metabolism were upregulated in a manner which is similar to the effect of MA-5 in NDUFS-4 knockout cochlear (Fig. 6d). These data suggest that similar transcriptional mechanisms are affected in MA-5-treated cochlea and HEI-OC1 cells, including the downregulation of EMT, hypoxia, TNF-α signaling NF-κB, and KRAS signaling and upregulation of heme metabolism, Myc targets and xenobiotic metabolism.

**Figure 6.**
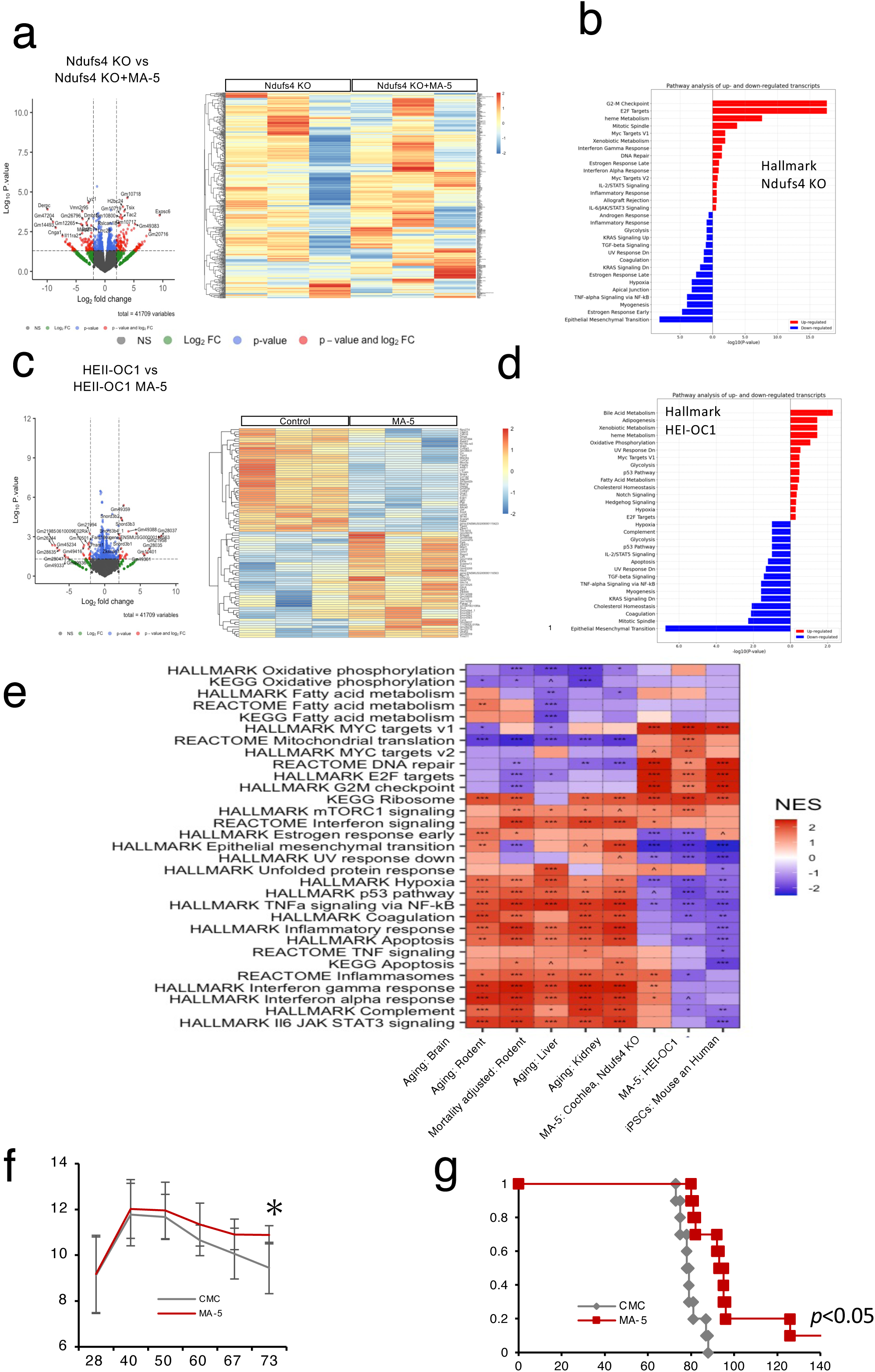
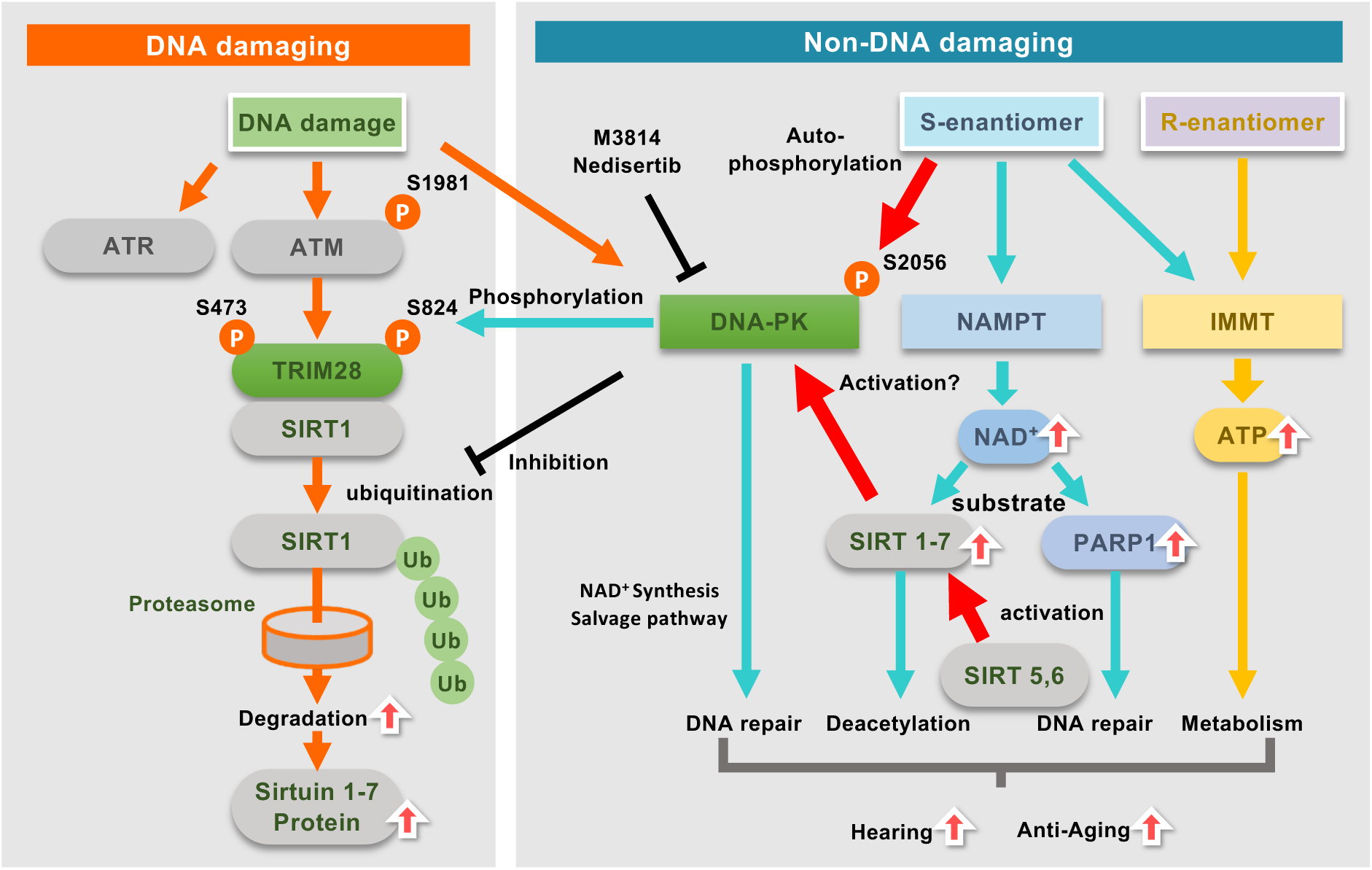
MA-5 follows a metabolic pathway similar to that of iPS cells.” **a**, Volcano plots of WT and Ndufs4 KO cochlear cells treated with MA-5. The x-axis represents the log2 fold change in gene expression, and the y-axis represents the -log10 P-value. Red dots indicate genes that are significantly differentially expressed in both p-value and logFC. Total = 41,709 variables. Heatmap of DEG Z- scores from cochlea. Rows represent individual genes. Columns show WT and Ndufs4 KO conditions. differentially expressed genes (DEGs) were filtered using *p* < 0.05, Max1, and SD1. **b**, Pathway analysis of DEGs in cochlear cells treated with MA-5. Up-regulated pathways identified by KEGG analysis are shown in red, and down-regulated pathways are shown in blue. **c**, Volcano plots of HEI-OC1and HEI-OC1 cells treated with MA-5. Pathway analysis of DEGs in HEI-OC1 cells treated with MA-5. **d**, Pathway analysis of DEGs in HEI-OC1 cells treated with MA-5. Up-regulated pathways identified by KEGG analysis are shown in red, and down-regulated pathways are shown in blue. (C) WT cochlear cells compared to WT control. **e**, Enrichment of pathways by MA-5 treatment and aging. Normalized Enrichment Score (NES) 0.05 < ^p-adj < 0.1; *p-adj < 0.05; **p-adj < 0.01; ***p-adj < 0.001. Benjamini-Hochberg approach. **f**, Body weight of NDUFS4 KO mice treated with MA-5 (10 mg/kg). Data were mean ± SE. \**p*<0.05, \*\**p*<0.01 vs. DMSO control; statistics were calculated by unpaired *t* test. **g**, Survival curve of NDUFS4 KO mice treated with MA-5 (10 mg/kg). Data were mean ± SE. \**p*<0.05, \*\**p*<0.01 vs. DMSO control; statistics were calculated by Kaplan-Meier method. **h**, Graphic summary. Triad mechanism of S-enantiomer; 1) ATP synthesis by binding with IMMT; 2) NAD synthesis by activating NAMPT; and 3) SIRT upregulation through TRIM28 phosphorylation by DNA-PK activation.

Aging is a time-dependent decline in functional capacity across the lifespan, characterized by the accumulation of molecular damage resulting from a diminished damage repair capacity^52^. The epigenome is a dynamic maintenance system that regulates chromatin organization and gene activity through various chemical modifications without altering the DNA sequences. However, changes to the epigenetic clock typically occur after several years of drug administration^53^. Recently, RNA-seq was found to characterize the gene expression signatures of mammalian longevity and identified tissue-specific and universal transcriptomic biomarkers of mammalian aging and mortality^54,55^. Kriukov et al. revealed that, in the iPS signature, EMT, protein secretion, TNF-α signaling via NFkB, hypoxia, UV response, hypoxia, apical junction, unfolded protein response, myogenesis, P53 pathway, apoptosis, inflammatory response, and IL6 JAK STAT3 signaling were downregulated, while E2F targets, G2M, checkpoint, Myc targets, DNA repair, ribosome, mitochondrial translation and the ATP metabolic process were upregulated by the Yamanaka factor (OSKM) introduction^56^. In addition, Yang et al. also reported that EMT, apoptosis, inflammatory response, TNF-a signaling via NF-kB, IL-6, JAK-STAT signaling, and coagulation were downregulated, while DNA repair, base excision repair, extension of telomeres, and Myc targets were upregulated by OSK^55^. Therefore, we conducted a gene set enrichment analysis (GSEA) to identify the pathways responsible for the similarities and differences between the transcriptomic signatures of MA-5, aging, mortality, and OSKM-related iPSCs. This analysis utilized the KEGG gene database, hallmark gene sets, and Reactome pathway database^54^ (Figure 6e). When the iPSC signatures were compared, MA-5 showed similar effects by downregulating EMT, UV response down, hypoxia, p53 pathway, TNF-α signaling via NF-κb, coagulation, apoptosis, and complement, and by upregulating Myc targets, DNA repair, E2F targets and ribosome. They are positively correlated with iPSC populations and negatively associated with mammalian age-related changes occurring in specific organs, such as the brain and liver, as well as across multiple tissues of mice, rats, and humans. Despite the differences in some pathways, MA-5 predominantly affected the same group of genes as epigenetic reprogramming by OSK, suggesting that these effects may be mediated through shared pathways.

### MA-5 improved NDUFS4 knockout mouse survival

Finally, to investigate whether MA-5 contributes to a reduction of mitochondrial disease model life span, we orally administered 10 mg/kg to NDUFS4 KO mice until their end of life. MA-5-fed mice showed high body weight compared with control (Fig. 6f) and l significantly prolonged survival compared to control mice (*p*<0.05, Fig. 6g). Collectively, these findings suggest that MA-5 functions as a modulator of aging in mice with mitochondrial dysfunction, as its administration leads to increased lifespan under mitochondrial damaging condition.

## Discussion

### Synergistic activation of NAD^+^/SIRT system by S-enantiomer

A major unsolved problem in aging research is the development of drugs for longevity interventions. We report here that S-enantiomer of exhibited three distinct metabolic pathways: 1) ATP synthesis by binding with IMMT; 2) NAD synthesis by activating NAMPT; and 3) SIRT upregulation through TRIM28 phosphorylation by DNA-PK activation (Fig. 6h, Graphic Summary). Notably, activation of these three pathways is not associated with DNA damage; rather, it facilitates DNA repair. The NAD^+^/SIRT system plays a crucial role in cellular energy metabolism, hearing, aging, and cell stress response^57^. So far, the conventional methods for modulating the NAD^+^/SIRT system include, 1. the oral administration of NAD^+^ precursors (NAM and NMN), 2. NAD^+^ synthesis activators (NAMPT activators and ACMSD inhibitors) or NAD^+^ consumer inhibitors (PARP and CD38 inhibitors)^58^, and 3. SIRT activators. Concerning oral supplementation of NAD^+^ precursors, the clinical effects of orally administered NAM and NMN remain an area of ongoing research but high-level NAM administration can exert negative effects through multiple routes^59^. Currently, Several NAMPT activators have been reported^60^. SBI-797812 activated NAMPT-mediated NMN production in the presence of NAM, PRPP, and ATP. The ability of SBI-797812 to activate NAMPT was abolished by the NAMPT inhibitor, FK-866 by binding to the NAMPT active site^60^. Our docking simulation suggested that S-enantiomer binds to the active site of NAMPT, which overlaps with the binding site of FK866, and that given that the structure of S-enantiomer is smaller than FK866, it is likely that S-enantiomer enhances the activity of NAMPT by avoiding steric hindrance with its substrate. In addition, because NMPT activity is highly dependent on the ATP concentration^61^, the synergistic effect of S- enantiomer that increases the ATP and NAD^+^ is beneficial for activating the NAMPT activity. Because the two NAD^+^-consuming enzymes are SIRTs and PARPs, PARP1 inhibitors have been also reported to increase NAD^+^ levels^62^. PARP1 inhibitor improves age-induced cell damage and increasing the SIRT1 activity^63^. However, since PARP activity acts as a “guardian of the genome”^64^, inhibiting PARP1 may not be a beneficial strategy for improving damaged cell function. SIRT activators (resveratrol, SRT1720, MDL-800 etc.) have been also reported. Resveratrol can promote SIRT1 activity, but it remains unclear whether its effects are direct or indirect^65^. SRT1720 also exhibited off-target effects on non-SIRT proteins, including AMPK^66^. Consequently, the application of SIRT activators remains confined to basic research, with a significant gap between their potential and clinical practice^67^. On the other hand, MA-5 is currently undergoing phase II clinical trials in Japan, and in that sense, it can be considered a compound approaching clinical application.

### TRIM28 activation and DNA-PK activation

As mentioned above, various studies have been conducted on methods to increase NAD^+^ and methods to activate SIRTs, but no method to increase SIRT protein has been explored. In addition, increased NAD^+^ levels do not promote SIRT1 expression^28^. We found that S-enantiomer enhanced the phosphorylation of TRIM28, suppressing the ubiquitination of SIRT1 and increasing the SIRT1-7 protein expression levels. To date, the phosphorylation of TRIM28 has generally been considered a consequence of DNA damage^44^. However, we revealed that S- enantiomer does not evoke DNA damage, but rather promotes DNA repair, suggesting that TRIM28 phosphorylation by S-enantiomer is a new approach to modulate sirtuin proteins. During our investigation into the mechanism by which the S-enantiomer increases SIRT protein levels, we found that the S-enantiomer induces phosphorylation of DNA-PK, which subsequently phosphorylates TRIM28. Under DNA damage, ATM, ATR, and DNA-PK coordinate to facilitate DNA repair^44^. Among these, DNA-PK is a key enzyme that regulates the non-homologous end-joining pathway, which repairs DNA double-strand breaks resulting from DNA damage. However, our findings indicate that the S-enantiomer does not induce DNA damage but instead promotes DNA repair. Furthermore, the activation of DNA-PK by the S-enantiomer is not a consequence of DNA damage but is mediated through the phosphorylation of DNA-PKcs. Although we performed preliminary simulation (Extended Data Fig. 7), the precise binding site of the S-enantiomer and the detailed mechanism underlying its autophosphorylation remain to be elucidated in future studies.

Recent studies have revealed that DNA-PKcs is involved not only in canonical DNA repair but also in the regulation of transcription, telomere maintenance, metabolic processes, and mitochondrial function ^68^. In this context, the activation of DNA-PKcs by the S-enantiomer may have therapeutic potential not only for DNA damage repair but also for various aging-related diseases. Moreover, because DNA-PK activity is ATP-dependent, the ability of MA-5 and the S-enantiomer to enhance intracellular ATP levels is likely advantageous for DNA-PK activation. It is well known that all SIRT1-7 play important roles in the DNA damage response and aging. Especially, SIRT2^69^, SIRT5^70^ and SIRT6^71^ acetylate DNA-PK and activate its function. Furthermore, SIRT1^72^ and SIRT6^72^ interact with TRIM28, while SIR6 interacts with PARP1. These multiple interactions between SIRTs, DNA-PK, PARP1, and TRIM28 induced by S-enantiomer contribute to improving hearing cell survival and prolonged survival of NDUFS-4 KO mice.

DNA damage is a central cause of aging and failure to repair DNA leads to cellular senescence, apoptosis, or dysfunction, which accelerates aging^73^. DNA damage also contributes to chronic inflammation in aging through cGAS-STING and NF-κB pathways^74^. The triggers for activation of these signaling cascades are also the age-related accumulation of DNA damage. Because DNA-PK plays a central role in DNA repair, upregulation of DNA-PK function may contribute to the maintenance of genomic stability and the modulation of aging. On the other hand, Park et al. reported that DNA-PK negatively regulates AMPK, contributing to metabolic and fitness during aging^75^. S-enantiomer is a novel compound that not only enhances mitochondrial function, but also increases NAD⁺ levels, activates SIRT1 and PARP1, and upregulates SIRT1–7 proteins with activating DNA-PK phosphorylation. These multifaceted actions suggest that it exert effects distinct from those of conventional DNA-PK activators, as evidenced by its ability to extend the lifespan in mouse models.

### MA-5 mimics the pluripotency signatures

We performed GSEA comparison to identify the pathways responsible for the similarities and differences between the chemical treatments and OSK(M)-related iPSCs and the signatures of aging^55^. The anti-aging effects of chemical treatments were associated with the upregulation of respiration-associated pathways and the downregulation of hypoxia and multiple inflammation-associated pathways^55,56^. In the iPSC population, significant downregulation of EMT and apoptosis^76^ were observed. In addition, mitochondrial function, E2F targets, GM2 checkpoint, Myc target V1, base excision repair, DNA repair, telomere extension, and ribosomes are also upregulated in iPSCss^55,56^. Because MA-5 effectively counteracted these molecular hallmarks of aging, these data indicated the potential impact on aging and EMT-related diseases^77^. These findings further indicated that MA-5 affects a specific cellular process associated with longevity and rejuvenation, which is similar to reprogramming. In addition to its role in SIRT1 ubiquitination, TRIM28 has been reported to maintain the undifferentiated stem cell state^78^. In ES cells, the phosphorylation of TRIM28 at Ser824 causes chromatin relaxation^79^, which maintains the undifferentiated state and the expression of ES cell-specific markers such as Oct4 and SSEA^78^. In this regard, TRIM28 phosphorylation may be also an important factor that controls the reprogramming efficiency of stem cells and the quality of iPS cells^78^. Additionally, direct interaction between OCT4 and DNA-PKcs and identified specific OCT4 and DNA-PKcs binding sites and DNA-PKcs as an upstream mediator of OCT4-induced MYC activation has reported^80^. The dual stimulation of increased NAD^+^production and upregulation of SIRT1-7 protein expression may further support this role.

### In conclusion

The S-enantiomer of tMA-5 belongs to a novel class of compounds that modulate fundamental pathways associated with lifespan extension, including ATP and NAD⁺ production, upregulation of SIRT proteins, and activation of DNA-PK. Therefore, it holds strong potential as an oral therapeutic agent for the regulation of aging

## Abbreviations

ABR: auditory brainstem response
AMPK: adenosine monophosphate-activated protein kinase
APAP: acetaminophen
ARHL: age-related hearing loss
ATM: Ataxia telangiectasia mutated
ATP: adenosine triphosphate
ATR: Rad3-related ATR
BSO: l-buthionine-(S, R)-sulfoximine
CPT: camptothecin
DEG: differentially expressed genes
DIHL: drug-induced hearing loss
DNA-PK: DNA-dependent protein kinase
EMT: epithelial mesenchymal transition
GSEA: gene set enrichment analysis
HEI-OC1: House Ear Institute Organ of Corti 1
IHC: inner hair cell
NAD^+^: nicotinamide adenine dinucleotide
NMNAT: NAM mononucleotide adenylyl transferase
NAMPT: nicotinamide phosphoribosyltransferase
Ndufs4: NADH dehydrogenase (ubiquinone) Fe-S protein 4
NHEJ: non-homologous end joining.
NIHL: noise-induced hearing loss
PARP: poly(ADP-ribose) polymerase
SIRT: sirtuins, SGC; spiral ganglion cell
TRIM28: tripartite motif containing 28

## ACKNOWLEDGEMENT

We thank Maiko Ueda (Histological platform, Tohoku University School of Medicine) for their histological assistance. We also acknowledge the technical assistance of the Biomedical Research Core of the Tohoku University Graduate School of Medicine and the Biomedical Research Unit of Tohoku University Hospital.

## DATAAVAILABILITY

All mouse sequencing generated for this project are are deposited as PRJNA1177978 and available.

## AUTHOR CONTRIBUTIONS

Y.H., T.S., R.K. S.K and T.A. participated in the conception and design of the study, K.H. synthesized MA-5, R.K. and Y.M. synthesized enantiomers, R.K., Y.M. and R.U., D. S. prepared and measured the MA-5 and enantiomers concentration, Y.T. generated recombinant protein, Y.T., H.K., S. N., K.T. prepared protein simulation and modelling, H.K. performed NAMPT binding simulation assay and Biolayer Interferometry binding assay, A.T., Z.B., T.K. and Y.T. performed RNA-Seq data analysis, S.C. and Y.T. generated TRIM28 KO mouse and TRIM28 antibody, C.S. and M.F. generated human iPS-derived ear cells, T.M., Y.O., H. K. and K.M. collected human skin fibroblasts and bioenergetic assay, T.S. carried out RNA sequencing and analysis of cochlea, F.N. examined NDUFS4 KO survival examination, K. T. performed cochlear imaging, D.S. measured MA-5 concentration in the cochlea, Y.H., J.S., and C.S. performed animal experiment and measured hearing capacity, T.S. performed DNA damage imaging, Y.O., K.N., H.E., Y.K., K.H., T.T., K. N., H. E., P.A., V.N.D.Y.K. and Y.T. discussed the results, Y.H. and T.A. conceived and designed this research; K.N., H.E. and T.A for funding acquisition, T.A. conceptualization; funding acquisition; supervision; writing – review and editing.

## METHODS

### Animals and experimental protocol

Male and female wild-type (WT) and Ndufs4 KO mice with C57BL/6J genetic background (B6.129S4-Ndufs4tm1.1Rpa/J) were used in this study (4 weeks of age, weighing 8-15 g at study onset). Because our preliminary experiments showed that Ndufs4 KO mice exhibit early onset progressive hearing loss as early as 6 weeks of age and premature death around 7 weeks of age, we decided to use mice aged 4 to 6 weeks for these experiments to avoid the effect of progressive hearing loss. Heterozygous Ndufs4 KO mice were crossed, and their offspring were genotyped through polymerase chain reaction (PCR) with the following primers to obtain WT and homozygous Ndufs4 KO mice. All mice were treated following the guidelines presented in the Standards for Human Care and Use of Laboratory Animals of Tohoku University, Guidelines for Proper Conduct of Animal Experiments by the Ministry of Education, Culture, Sports, Science, and Technology of Japan, and the National Institute of Health Guide for the Care and Use of Laboratory Animals. All the animal experiments were approved by the Ethics Committee for Animal Experiments of Experiments of Tohoku University Graduate School of Medicine (2019-249). The mice were divided into three experimental groups. In group 1, seven male WT mice and seven male Ndufs4 KO mice were assigned and sampled on postnatal (P) day 32 (P32). Three mice were used for surface preparations and paraffin sections in both WT and Ndufs4 KO groups, and four mice were used for RNA sequencing. In group 2, five female WT mice and five female Ndufs4 KO mice were assigned and subjected to 100 dB acoustic exposure on P34. Subsequently, ABR thresholds were measured 2 days before and 1 and 8 days after acoustic exposure. After the sampling on P42, the left cochlea was used for surface preparation, and the right was used for paraffin sections. In group 3, six male WT mice and six male Ndufs4 KO mice were assigned and subjected to 86 dB acoustic exposure on P34. Afterward, the same process as described above was performed. These mice were genotyped at P-21. Both male and female mice were used in the study. The survival curves were analyzed by pooling the data from both sexes (male-to-female ratio is ∼1:1), as there was no sex-specific difference in survival ship. Survival of KO mice was recorded.

### Hearing function test

The ABR threshold was measured as described previously^81^. The mice were anesthetized using ketamine (100 mg/kg body weight) and xylazine (20 mg/kg body weight) through intraperitoneal administration. Needle electrodes were placed subcutaneously at the vertex (active electrode), base of the ear pinna (reference electrode), and back (ground electrode). ABR recordings were performed using a TDT System 3 auditory-evoked potential workstation and BioSigRP software (Tucker-Davis Technologies, Alachua, FL, USA). ABR responses were evoked using bursts of pure tones at frequencies of 4, 8, 12, 16, and 32 kHz. Evoked responses were averaged across 1,000 sweeps. Responses were collected for stimulus levels in 5 dB steps from 100 dB sound pressure level (SPL) to 10 dB SPL. The measurement of ABR and determination of the threshold were performed by separate examiners^15^. Trim28 heterozygote mice and wild type littermate were kindly provided by Prof. Shunsuke Chikuma (Department of Microbiology and Immunology, Keio University School of Medicine, Tokyo). TRIM28 Flox mice (Cammas et al. https://pubmed.ncbi.nlm.nih.gov/10851139/) were kindly provided by Drs. Pierre Chambon and Regine Losson), backcrossed into C57BL/6 background for 8 times (https://pubmed.ncbi.nlm.nih.gov/22544392/, https://pubmed.ncbi.nlm.nih.gov/33619215/) and maintained at a SPF facility of Keio University. TRIM28 haplo-insufficient mice (heterozygote: hereafter Trin28+/-) are obtained from a breeding colony of Female TRIM28 Flox mice and x male B6(Cg)-*Foxn1^tm3(cre)Nrm^*/J mice. The male B6(Cg)-*Foxn1^tm3(cre)Nrm^*/J express CRE on the germ cell, worked as a “CRE deleter” (https://pubmed.ncbi.nlm.nih.gov/27880796/). The offspring from FoxN1-CRE x Trim28 lox carried germline Trin28+/- allele and used to generate Trin28+/- mice by breeding with wild-type C57BL/6 mice.

### Human skin fibroblasts

Fibroblasts obtained by skin biopsy of mitochondrial disease patients were collected in Chiba Children’s Medical Center and Tohoku University Hospital under the approval of the Ethical Committee of Tohoku University. Written informed consent was obtained from all the patients. NDUFS-4 and MELAS are examined as previous reported^11^. In the cell viability experiments, fibroblasts were cultured in 1.0 g/L low-glucose DMEM with 10% FBS at 37°C in 5% CO_2_. Cell viability assay was performed ^11^. The LDH level was measured by cell count using LDH cytotoxicity detection kit (Takara).

### Cell culture and viability, ATP and NAD^+^ assays

Fibroblasts obtained by skin biopsy of mitochondrial disease and congenital hearing loss patients were collected in Kanagawa Children’s Medical Center (KCMC), Jichi Medical University and Tohoku University Hospital under the approval of the Ethical Committee of Tohoku University (2021-1-901, 2024-1-905). Written informed consent was obtained from all the patients. Fibroblasts from patient with mitochondrial diseases, congenital hearing loss, Trim28 heterozygote mice and their wild type littermates were cultured in 1.0 g/L low glucose DMEM with 10% FBS, 100 units/ml penicillin and 100 μg/ml streptomycin (Normal growth medium) at 37°C, 95% ambient air and 5% CO_2_ and 21% oxygen for maintaining culture. In cell viability assay, low glucose DMEM with 1% FBS and antibiotics (assay medium) was used for fibroblasts. HEI-OC1 cells^22,23^ were provided from Kalinec, GM (UCLA) and maintained in 4.5 g/L high-glucose DMEM containing10% FBS at 33 ^◦^C,95% ambient air and 5% CO_2_ and 21% oxygen for maintaining culture. In cell viability assay, low glucose DMEM with 5% FBS (assay medium) was used for HEI-OC1 cells. In the cell viability assay, skin fibroblasts and HEI-OC1 cells were plated in 96 well plates *at a density of* 3 x 10^3^ cells /100*μl* assay medium per each well. After 24 h of incubation, L-buthionine-(*S*,*R*)-sulfoximine (BSO) cisplatin, APAP and MA-5 or 0.1% DMSO was added by assay medium exchange with compounds. After each compound was applied at mentioned final concentrations, fibroblasts and HEI-OC1 cells were cultured for 48hr and cell viability was measured by Cell count Regent SF (nacalai tesque) and the cellular damage was measured by LDH level in culture medium using LDH cytotoxicity detection kit (Takara) according to manufacture instruction. In ATP and NAD^+^ assays, skin fibroblasts *at a density of* 1.5x 10^3^cells /100*μl* assay medium and HEI-OC1 cells *at a density of* 6x 10^3^ cells /100*μl* assay mediums per each well were plated in 96 well plates. After 24 h of incubation, various concentration of MA-5 racemate, MA-5 S-enantiomer and R-enantiomer were added by assay medium exchange with compounds. The ATP level and the NAD^+^ level was measured after 3 hrs and 6hrs from compounds treatment by ’’Cell’’ ATP Assay reagent Ver.2 (Toyo ink, Osaka, Japan) and NAD+/NADH Glo Assay kit (Promega) according to manufacture instruction.

### Organ of Corti explants Organotypic cultures^82^

Postnatal day (P) 3 C57BL/6 mice were used in our experiments. Organ of Corti explants from P3 C57BL/6 mice was dissected and placed in culture dishes. Explant incubation was performed in DMEM/F12 containing 1× N2 (100×, Invitrogen, 17502-048) and 1× B27 (50×, Invitrogen, 17504-044) in a humid environment with 5% CO2 at 37°C. For the assessment of toxicity of cisplatin, the Organ of Corti explants were pre-treated with 10 μM of MA-5 or vehicle control for 2hrs and cotreated with varying concentrations (0, 5, 7. 5, 10, 15, 20 μM) of cisplatin and 10 μM of MA-5 or vehicle control for 24hr. For the experiments of cell protective effects of MA-5 against cisplatin, the Organ of Corti explants were pretreatment with varying concentrations (0, 1, 5, 10, 50 μM) of MA-5 for 2 hr and then cotreated with 10 μM of cisplatin (Cis + MA-5) for 24 hr.

### Human cochlear epithelial cells derived from induced pluripotent stem cells (iPSCs)

Human iPSCs were cultured and differentiated into otic progenitor cells-like cells (OPCs) as described previously^83^. OPCs were dissociated with Accutase (nacalai tesque) and cultured in 96-well ultra-low attachment plates (Corning) to form spheres in DMEM/F12 (FUJIFILM Wako Pure Chemical) containing 1% N2 supplement (Thermo Fischer Scientific), 2% B27-vitamin A supplement (Thermo Fischer Scientific), 2 mM Glutamine (nacalai tesque), 10 ng/mL FGF2 (R&D systems), 20 ng/mL EGF (Pepro Tech), 50 ng/mL IGF-1(R&D systems), 3 mM CHIR99021(Focus Biomolecules), 0.5 mM retinoic acid (Sigma), 20 ng/mL Wnt3a (R&D systems), 50 ng/mL heparan sulfate (Sigma), and 10 mM Y27632 (FUJIFILM Wako Pure Chemical). On day 5, the medium was changed once. On day 7, the cells were harvested, dissociated with Accutase, frozen in STEMCELL BANKER (Takara), and stored in LqN2 until use. To differentiate inner ear-like cells expressing POU4F3 and MYO7A, the cells were thawed and plated on Matrigel (Corning, #354277)-coated plates. They were cultured in DMEM/F12 with 1% N2 supplement, 2% B27-vitamin A supplement, 2 mM Glutamine,10 ng/mL FGF2, 20 ng/mL EGF, 50 ng/mL IGF-1, 3 mM CHIR99021, 0.5 mM retinoic acid, and 10mM Y27632 (monolayer day 1). The medium was changed on monolayer days 2 and day 5 to DMEM/F12 with 1% N2, 2% B27-vitamin A, 2 mM Glutamine, 20 ng/mL EGF, 10 ng/mL IGF-1, and 3 mM CHIR99021. Cells were harvested and analyzed on monolayer day 8.

### Measurement of MA-5 concentration in the cochlear

Chemical standard of MA-4, mono-fluoro type analogue of MA-5 IUPAC name 4-(4-fluorophenyl)-2-(1H-indole-3-yl)-4-oxobutanoic acid was synthesized as an internal standard (IS). The LC-MS/MS analysis was performed on NANOSPACE SI-II system equipped with a dual pump, an autosampler, a column oven (Osaka Soda, Osaka, Japan) interfaced to a TSQ quantum ultra, MS/MS system equipped with ESI operated in positive ion mode (Thermo Fisher Scientific, Waltham, MA, USA). The MS/MS was performed using the multiple reaction monitoring (MRM) mode. LC separation was performed using a reverse-phase column (Xbridge C18 100 mm × 2.0 mm i.d., 3.5 µm particle size; Waters Corp., Milford, MA, USA) with a gradient elution of solvent

A (5 mmol/L ammonium acetate in water) and solvent B (5 mmol/L ammonium acetate in water/methanol, 5/95, v/v%) at 0.2 mL/min. The oven temperature was 40°C. The data were collected using the Xcalibur software (Thermo Fisher Scientific) and the ratio of the peak area of analyte to the IS was regulated with the calibration curve made by the standard solution.

### Enantiomer separation and measurement

The S-enantiomer and R-enantiomer of MA-5 were optically prepared from racemic MA-5 using a LC-Forte preparative system (YMC, Kyoto, Japan) equipped with a CHIRALPAK AD-H column (20 mm i.d. × 2500 mm, 5 µm particle size; DAICEL, Osaka, Japan). The crude powder was purified by silica gel column chromatography and recrystallization to afford the enantiopure MA-5 enantiomers. Each enantiomer of MA-5 was dissolved in NaOH (0.1 mol/L)–saline (1:9) and administered intravenously to a male rhesus monkey at a dose of 1 mg/kg. Plasma samples were collected sequentially from 5 minutes to 24 hours after administration, and the concentration of each MA-5 enantiomer in the plasma samples was quantified using the following method. The plasma sample from a rhesus monkey was deproteinized with acetonitrile–formic acid (100:0.1, v/v) containing an internal standard MA-5-d6 (1 µg/mL, final concentration), and 2 µL of the supernatant was subjected to LC-MS/MS analysis. LC–MS/MS analysis was performed on an NANOSPACE SI-2 system (Osaka Soda, Osaka, Japan) coupled to a TSQ Quantum Ultra triple quadrupole mass spectrometer (Thermo Fisher Scientific, Waltham, MA, USA). Chromatographic separation was performed on a CHIRALPAK AD-3R column (2.1 mm i.d. × 150 mm, 3 µm; DAICEL, Osaka, Japan) at 35 °C. Mobile phase was ddH2O–formic acid (100:0.1, v/v) for phase A and acetonitrile–formic acid (100:0.1, v/v) for phase B, and the analytes were eluted isocratically at 43% phase B for 9.7 min. S-enantiomer was eluted in 6.9 min, while R-enantiomer was eluted in 8.7 min. The analytes were ionized using the negative ion mode ESI probe and detected in MRM mode. The mass transition conditions were m/z 328.1–116.0 for each enantiomer of MA-5 and m/z 334.1–122.0 for each enantiomer of MA-5-d6. LC– MS/MS control, data acquisition, and data processing were performed using Xcalibur (Thermo Fisher Scientific).

### siRNA Knock down

siRNA knock down of TRIM28, ATM and DNA-PKcs were performed. Briefly, Silencer® Select Pre-Designed siRNA of TRIM28, ATM and DNA-PKcs or Silencer™ Select Negative Control No. 1 siRNA was transfected in HEI-OC1 cells with Liofectamine RNAiMAX and Opti-MEM I Reduced Serum medium by reverse transfection method. 48 hr after siRNA transfection, Vehicle or MA-5 S-enantiomer was administrated and incubated another 24hr. Protein samples were extracted from cells for Western blot analysis as mentioned.

### SIRT1 protein poly-ubiquitination analysis

For Myc-tag pull down for SIRT1 protein poly-ubiquitination analysis, over expression of Myc-tagged ubiquitin and Flag-tagged SIRT1 protein DNA expression vectors were used. HEI-OC1 cells were cultured in 10cm cell culture plate at 2 x 10^4^ cells and incubated for 24hrs. 6μg of DNA-expression vector, 25μl of P3000 regent and 25μl of Lipofectamine 3000 in 1.2ml of Opti-MEM I Reduced Serum medium were prepared as manufacture instruction and applied in HEI-OC1 culture in 10cm for transfection. 24hr after transfection started, vehicle (DMSO) or MA-5 S-enantiomer were added and incubated for another 24hr. Protein samples were extracted from cells for Western blot analysis as mentioned.

### Real time Quantitative PCR

HEI-OC1 cells were homogenized in Sepazol-RNA I Super G (Nacalai Tesque, Inc., Kyoto, Japan), and total RNA was extracted according to the manufacturer’s instructions. cDNA synthesis was performed using ReverTra Ace qPCR RT Master Mix with gDNA Remover (Toyobo Co., Ltd. Osaka, Japan). Quantitative PCR was performed using the TaqMan RT-qPCR system (Applied Biosystems, Waltham, MA, USA). A thermal cycling condition used in this study is as follows; preheat at 95 °C for 1min, then 15 sec at 95 °C and 3 at 60 °C followed by 40 cycles. TaqMan Primers were purchased from Applied Bio systems/Thermo Fisher Scientific (Extended Table 1). The sequences of the primers and probes are certificated by the company, but not open based on the company policy.

### Cochleae Immunohistchemisty^82^

Cochleae fixed in 4% PFA were rinsed three times with PBS and permeabilized with 1% Triton X-100 in PBS for 30 min at room temperature. Cochlear explants were then blocked with 10% donkey serum in PBS for 1-hour, followed by incubation with the specific primary antibodies overnight at 4 ^◦^C. The following primary antibodies were used: parvalbumin (1:500, Abcam, ab32895) and cleaved caspase-3 (1:500, Cell Signaling, 9661). Then the sections were incubated with secondary antibodies overnight: Alexa488-conjugated donkey anti-mouse IgG (1:500, Invitrogen) and Alexa594-conjugated donkey anti-rabbit IgG (1:500, Invitrogen).

### Immunostaining for mitochondria imaging with SoRa

Fixed cochlea tissues were sectioned at 4μm interval. After deparaffinization, the sectioned tissues were incubated in TE buffer (pH 9.0) at high temperature and pressure using a pressure cooker for antigen retrieval and permeabilization. SIRT1were labeled with antibody (Cat. No. ab189494, Abcam), followed by Cy3-Donkey anti-rabbit IgG antibody (Cat. No. 711-165-152, Jackson ImmunoResearch Labs). Cochlea tissues were imaged and image processing was performed with BZ-X800 (Keyence, Osaka, Japan). Cochlea tissues were imaged using the Nikon Ti2 microscope partnered with the Yokogawa CSU-W1 SoRa system using Apo TIRF ×100 oil DICN2 as an objective and taken as a z-stack at 0.12 μm per z step. A total of 5 tubules per sample were imaged^35,84^. Imaging with SoRa and image processing was performed through the Nikon imaging center in Osaka University.

### Computational details

#### System preparation

The 3D structures of Mitochonic acid 5 (MA-5) in (*R*)- and (*S*)-stereoisomers under neutral conditions were constructed and optimized using Gaussview and Gaussian16 with density functional theory (DFT), employing the B3LYP/6-31G* basis set. The partial charges and parameters for these ligands were generated using the restrained electrostatic potential (RESP) method in conjunction with the general AMBER force field 2 (GAFF2)^85^, implemented through the antechamber module within the AmberTools21 package, as previously described^34^.The dimeric Nicotinamide phosphoribosyltransferase (NAMPT), obtained from the Protein Data Bank (PDB ID: 8DSE^61^), was stripped of bound small molecules and then supplemented with missing residues using the MODELLER software^86^executed in UCSF Chimera version 1.17^87^. Subsequently, this protein was protonated under neutral conditions using the PDB2PQR web server^88^.

#### Binding Site Prediction

MA-5 binding sites on the dimeric NAMPT were identified through BDK and a custom Python script. Both (*R*)- and (*S*)-stereoisomers of MA-5, along with NAMPT, underwent conversion to PDBQT format using the ADFR package. Docking was conducted utilizing AutoDock Vina 1.2.3^89^ within a 90 × 90 × 90-Å box over 100 iterations, with 40 poses per run. Subsequently, a total of 4,000 docking results underwent *k*-means clustering to assess potential binding sites. To further refine the findings, blind docking outcomes were subjected to focused docking techniques, aimed at confirming the preferential stereoisomer type of MA-5 for each predicted binding location within a 20 × 20 × 20-Å box.

#### Molecular Dynamics Simulation

The topology and coordination files for (*R*)- and (*S*)-MA-5 bound to the active site of NAMPT were generated using the tLEaP module and the ff19SB force fields^90^. Subsequently, the SANDER program minimized the complexes while restraining the protein backbone. The complexes were simulated under periodic boundary conditions (PBC) using the AMBER20 package program for 3 replications, each individually set by varying the velocities. Each system was solvated in a periodic box using the TIP3P water model with a distance of 12 Å from the protein surface and neutralized with sodium ions^91^. The system was heated to 300 K for 20 ps using the canonical ensemble (NVT). Selected systems were simulated for 500 ns to evaluate ligand-binding stability over the MD trajectory by measuring the distance between the center of mass of the ligand and the binding pocket of the NAMPT active site using the CPPTRAJ module^92^. Only systems demonstrating stable binding to the active site over the 500 ns-trajectories were analyzed in terms of ligand-binding patterns and susceptibility using the MM-GBSA method^93^.

### Dockings simulation using MOE

Docking simulation of S-enantiomer to DNA-PK was performed using molecular operations environment (MOE) system (MOLSIS, Inc.). A small compound structure of S-enantiomer and a protein structure of DNA-PK (PDB ID: 7OTY) was prepared using standard protocols in the general docking of MOE. After preparation, the docking simulation was started, and all docked poses were ranked by the London dG score.

### Western blotting

Blood inside organs were removed by rinsing tissues with PBS. Tissues were snap-frozen under liquid nitrogen. Tissues were homogenized in RIPA buffer (Sigma) with protease (Roche), deacetylase (nicotinamide, trichostatin A) and phosphatase inhibitors (Roche). Protein concentrations of samples were determined by Lowry assay and each amount of protein (30–50 ug per sample) were loaded for SDS-PAGE. Antibodies from the following companies were used for Western blot analysis: SIRT1 (1:1000 dilution, #9475, Cell Signaling), SIRT2 (1:1000 dilution, #12650, Cell Signaling), SIRT3 (1:1000 dilution, #5490, Cell Signaling), SIRT4 (1:1000 dilution, #69786, Cell Signaling), SIRT5(1:1000 dilution, #8782, Cell Signaling), SIRT6 (1:1000 dilution, #12486, Cell Signaling), SIRT7(1:1000 dilution, sc-365344,Santa Cruz), IMMT/Mitofilin (1:1000 dilution, 10179-1-AP, ProteinTech), p53 (1:1000 dilution, #2524, Cell Signaling), acertyl p53(K305) (1:1000 dilution, ab109396, abcam), Phospho-p53 (Ser15) (1:1000 dilution, #9284, Cell Signaling), TRIM28(KAP1)(1:10000 dilution, ab109287, abcam), 4H11 pTRIM28 (Ser 473) mouse mAb(1:1000 dilution, Gift from Dr .Shunsuke Chikuma), pTRIM28(KAP1) (Ser 824) (1:1000 dilution, ab133440, abcam), ATM (1:1000 dilution, #2873, Cell Signaling), pATM(Ser 1981)(1:1000 dilution, Invitrogen #14-9046-82), DNA-PKcs (1:1000 dilution, #4602, Cell Signaling), pDNA-PKcs(Ser 2056) (1:1000 dilution, ab18192, abcam), K48-linkage Specific Polyubiquitin (1:1000 dilution, #4289, Cell Signaling), FLAG® tag epitope(DYKDDDDK tag) (1:1000 dilution, F1804, Sigma Aldrich), MYC Tag (1:1000 dilution, #2278, Cell Signaling), PARP1 (1:1000 dilution, #9542, Cell Signaling), PARP1, acetyl Lys521 (1:1000 dilution, CSB-PA890185, CUSABIO), poly(ADP-ribose) polymer (pADPr) (1:1000 dilution, sc-56198, Santa Cruz), 53BP1 (1:1000 dilution, ab175933, Abcam), Phospho-53BP1 (Ser25) (1:1000 dilution, NB100-1803, Novus Biologicals), Acetyl-Histone H3 (Lys18) (1:1000 dilution, #9675, Cell Signaling), Phospho-Chk2 (Thr68) (1:1000 dilution, E-AB-20843, Elabscience), DNA PKcs (pS2056) (1:1000 dilution, ab124918, Abcam), GAPDH (1:1000 dilution, sc-32233, Santa Cruz), β-actin (1:500 dilution, sc-47778, Santa Cruz). Secondary antibodies used in the Western blot experiment were Goat anti-Mouse IgG (H+L) Secondary Antibody, HRP (1:500 dilution, #31430, Invitrogen) and Goat anti-Rabbit IgG (H+L) Secondary Antibody, HRP (1:1000 dilution, #31460, Invitrogen). Images Chemiluminescence was performed using a Chemi-Lumi One Super immunochemistry kit (Nacalai Tesque, Inc.) and images were captured using the ImageQuant LAS 4000mini biomolecular imager (Fujifilm, Tokyo, Japan). Protein band Bands intensities were analyzed using ImageQuant LAS 4000 mini software (GE Healthcare, Little Chalfont, UK). The protein abundance was analyzed by densitometry with ImageJ and normalized with β-actin or GAPDH.

### Enzymatic activity assay (SIRT1, NAMPT, NMNAT, PARP and DNA-PKcs)

Effects of enantiomer MA-5 R body on SIRT1 enzyme activity were measured by SIRT1/Sir2 Deacetylase Fluorometric Assay Kit (Cat # CY-1151v2, CYcLex, MBL-Life Science, Tokyo, Japan).

SIRT1 enzyme activity in nuclear fraction protein from HEI-OC1 cells were also measured by SIRT1 enzyme activity was measured by SIRT1/Sir2 Deacetylase Fluorometric Assay Kit. Briefly, HEI-OC1 cells were cultured in 100mm dish at 200 x 10^4^ cells and incubated for 24hrs. Nuclear fraction proteins were extracted from HEI-OC1 cells after 3hr DMSO or MA-5 treatment (10μM and 30 μM) and SIRT1 enzymatic activity of nuclear fraction proteins were measured according to manufacture instruction. Effects of MA-5 racemate, enantiomer MA-5 S and enantiomer MA-R S on nicotinamide phosphoribosyltransferase (NAMPT) enzymatic activity, nicotinamide mononucleotide adenylyltransferase (NMNAT) enzymatic activity and poly(ADP-r ibose) polymerase (PARP) activity were a measured by Universal Colorimetric PARP Assay Kit with Histone-Coated Strip Wells (R & D systems) NAMPT Inhibitor Screening Assay Kit (Cat #.71276-1,BSP-Bioscience), CycLex NMNAT Colormetoric assay kit (Cat. #CY1252. CYcLex, MBL-Life Science) and Universal Colorimetric PARP Assay Kit with Histone-Coated Strip Wells (Cat. # 4677-096-K, R & D systems) respectively according to manufacture instruction.

### Total RNA sequencing

For RNA extraction, the cochlear tissues were freeze-fractured and powdered. Next, RNA was extracted and purified using RNAiso Plus according to the manufacturer’s instructions. The integrity of the RNA was checked using Agilent RNA 6000 Nano Kit (Cat# 5067-1511; Agilent) on the Bioanalyzer (Agilent). The RNA Integrity Nunber (RIN) in all samples was 3.2 to 6.1, thus, it was considered that RNAs have fragmentation and total RNA sequencing is suitable in this study. Using 500ng of the RNAs of each sample, the libraries were created using NEBNext Ultra II RNA Library Prep Kit for Illumina and NEBNext rRNA Depletion Kit v2 (New England Biolabs, Cat# E7770S and E7400L) according to the manufacturer’s instructions, and the final PCR was performed as 15 thermal cycles. Concentrations and size distributions of the libraries were measured using an Agilent DNA 7500 kit (Cat. #5067-1506; Agilent) on Bioanalyzer. All samples were passed for analyses on NGS equipment. The libraries were pooled and the concentrations were adjusted to 1 nM. The pooled libraries were subjected to denaturation and neutralization. Subsequently, the libraries were diluted to 1.8 pM and then applied for an NGS run on the NextSeq 500 System (Illumina) with NextSeq500/550 v2.5 (75 Cycles) Kits (Illumina, Cat. #20024906). The sequencing was performed with paired-end reads of 36 bases. After the sequencing run, FASTQ files were exported and basic information of the NGS run data was checked on CLC Genomics Workbench 22.0 software (CLC, QIAGEN). In the results, the PHRED-score as a quality score of the reads over 20 was confirmed for 99% of all reads, indicating the success of the data acquisition in the NGS run. The read number was approximately 28 to 37 million per sample as paired-end reads.

### Transcriptomic signatures of aging, mortality, and lifespan

To identify transcriptomic signatures of chronological age, lifespan-adjusted age, expected mortality and maximum lifespan based on the aggregated meta-dataset of rodent relative gene expression, we utilized linear mixed-effect model with REML criterion via lmer function from lme4 and lmerTest R packages^94^. We considered change in log-expression as an outcome variable and difference in trait of interest as an independent variable, while tissue and source of data were implemented in the model as random terms, and sex was included as a fixed term. To examine genes associated with age-adjusted differences in expected mortality (in log scale), lifespan-adjusted age and maximum lifespan, we introduced difference in chronological age into a mixed-effect model as a separate fixed term covariate. Genes were considered statistically significant if their *p* value, adjusted with Benjamini-Hochberg (BH) method^95^, was lower than 0.05. Pairwise overlaps between transcriptomic signatures associated with different traits were assessed separately for up- and downregulated genes with Fisher’s exact test, and together with Pearson’s chi-square test.

### Statistics and reproducibility

GraphPad Prism 8 software (GraphPad Software, Inc.) was used to perform statistical analysis. Two-tailed Student’s t test was used to test differences between two groups and differences between multiple groups were analyzed by two-way ANOVA. Data are shown as mean ± SEM of biological triplicates. Details of the statistical analysis for each experiment was indicated in the relevant figure legends. For all statistical tests, *p* values < 0.05 were considered statistically significant. All experiments except RIME were repeated at least three times with similar results.

## Extended Figures

**Extended Figure 1 Auditory brainstem response (ABR) test of wild type (WT) and Ndufs4 KO mice a,** At P28, ABR threshold (4, 8, 12, 16 and 32kHz) was not changed between WT (n=5) and NDUFS4 KO mice. Then NDUFS4 KO mice was divided into 2 groups, (n=5 in each group) and administration of vehicle or MA-5 was started.

**b**, Comparison of Auditory brainstem response (ABR) test of Ndufs4 KO mice at postnatal day 28 (P28) and P40 (n=15). ABR thresholds at 4, 8, 12, 16 and 32kHz were measured at P28 and P40. At 40, the ABR threshold at 8 and 12kHz were significantly elevated compared with the ABR threshold at P28. Then NDUFS4 KO mice was divided into 3 groups, (n=5 each group) and started administration of vehicle or MA-5 (low dose1.0mg/kg and high dose 10mg/kg) for 3 weeks from P40 to P60. Ndufs4 KO mice a. \**p*<0.05, \*\**p*<0.01 vs. Ndusf4 KO mice at P28.; statistics were calculated by unpaired two-side t-test.

**Extended Data Fig. 2 MA-5 and each enantiomers exhibited no direct effect on SIRT1 enzyme activity. a**, SIRT1 deacetylase activity was measured in vitro in the presence of the drug, its S-enantiomer, and R-enantiomer at concentrations of 10, 30, and 100 μM (n=3-4). Data were mean ± SE. statistics were calculated by One way anova Durnett’s multiple comparison test.

**b**, DNA PARylation (left). Immunoblot of anti-Su lysine Ab (middle) and anti-acetylation of lysine(K) Ab (right). **c.** PARP1 enzyme activity *in vitro* with MA-5, S-enantiomer and R-enantiomer at 10,30 and 100 μM. (n=4). Data were mean ± SE.; statistics were calculated by one way anova Durnett’s multiple comparison test.

**Extended Data Fig. 3 Effect of MA-5 and each enantiomer onNMNAT1 and NMNAT3 enzyme activities** NAMNAT 1 and NAMTA3 enzymatic activities were measured by bio-luminescence assay incubated in control (0.1 % DMSO) or MA-5 at 1mM with or without mitofilin recombinant protein (final concentration at 1.6μg/ml) at 25 °C 40min in 5min intervals.

**Extended Data Fig. 4 Binding assay for NAMPT using Biolayer Interferometry (BLI) a,** Recombinant human NAMPT were purified.

**b.** Recombinant human NAMPT was mobilized on avidin tips, and the sensors were exposed to ligand solutions and investigated the binding of MA-5 and its enantiomers to NAMPT.

The binding curves obtained with increasing concentrations of MA-5 (Upper panel). Conversely, no binding affinity was observed for the R-enantiomer (lower panel).

**Extended Data Fig. 5 Compound C only increased the SIR3 protein expression but not other sirtuins.** The protein expression levels of SIRT1-SIRT7 in HEI-OC1 cells treated with MA-5 S-enantiomer at 0, 10 and 10 μM with or without AMPK inhibitor, Compound C, at 5 μM for 24hr.

**Extended Data Fig. 6 Effect of E3 Ubiquitin ligase inhibitors on the SIRT1-7 prortein expressions.** The protein expression levels of SIRT1-SIRT7 in HEI-OC1 cells treated with 0.1 % DMSO vehicle control or MA-5 S-enantiomer at 30 μM with or without E3 Ubiquitin ligase inhibitors, Thalidomide (T), Lenalidomide (L), Pomalidomide (P) and NSC 66811(N) at 10μM for 24hr.

**Extended Data Fig. 7 Predicted binding mode of S-enantiomer to DNA-PK using MOE.** a, Left panel, predicted binding mode of S-enantiomer to DNA-PKcs (PDB ID: 7OTY). Right panel, predicted complex structure of S-enantiomer to DNA-PK.

(a) Left panel, binding mode of M3814 to DNA-PKcs. Right panel, Complex structure of M3814 to DNA-PKcs.

**Extended Data Figure 8 Relation between SIRT2, 5,6 and other sirtuins.** RNA expression level of SIRT2 (**a**), SIRT5 (**b**) and SIRT6 (**c**) siRNA knockdown in HEI-OC1 cells (n=3). Data were mean ± SEM. \**p*<0.05, \*\**p*<0.01 vs. DMSO control; ^#^*p*<0.05, ^##^*p*<0.01 vs. negative control (NC); statistics were calculated by unpaired two-side t-test.

## Supplementary Table 1

(a) Taqman primer list used.

(b) siRNA list used.

