## Supplementary Fig.1-8+Table 1 for "Non-DNA-damaging DNA-PK activation improving hearing and prolonging life due to NAD^+^ and SIRT upregulation"

### Extended Data Figure 1

a

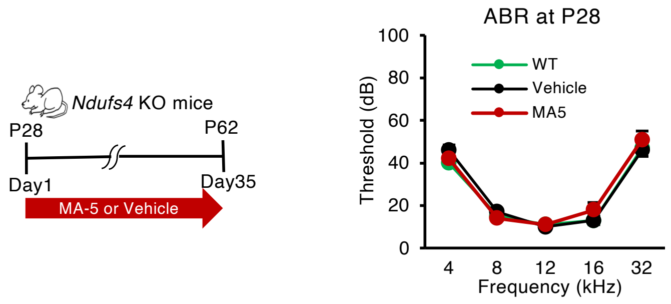


b

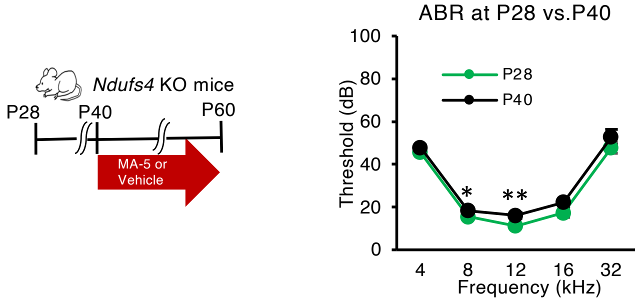

### Extended Data Figure 2

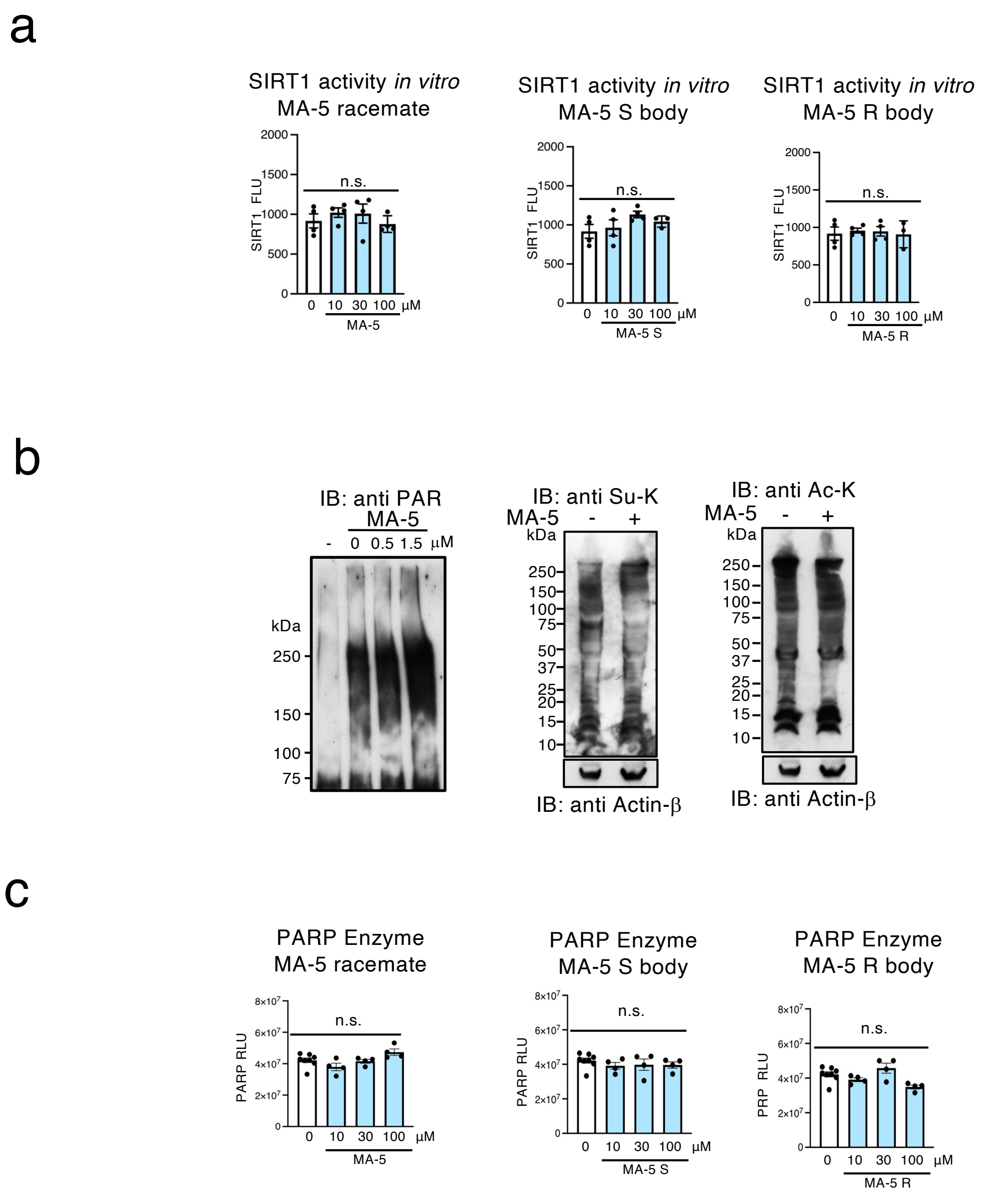

### Extended Data Figure 3

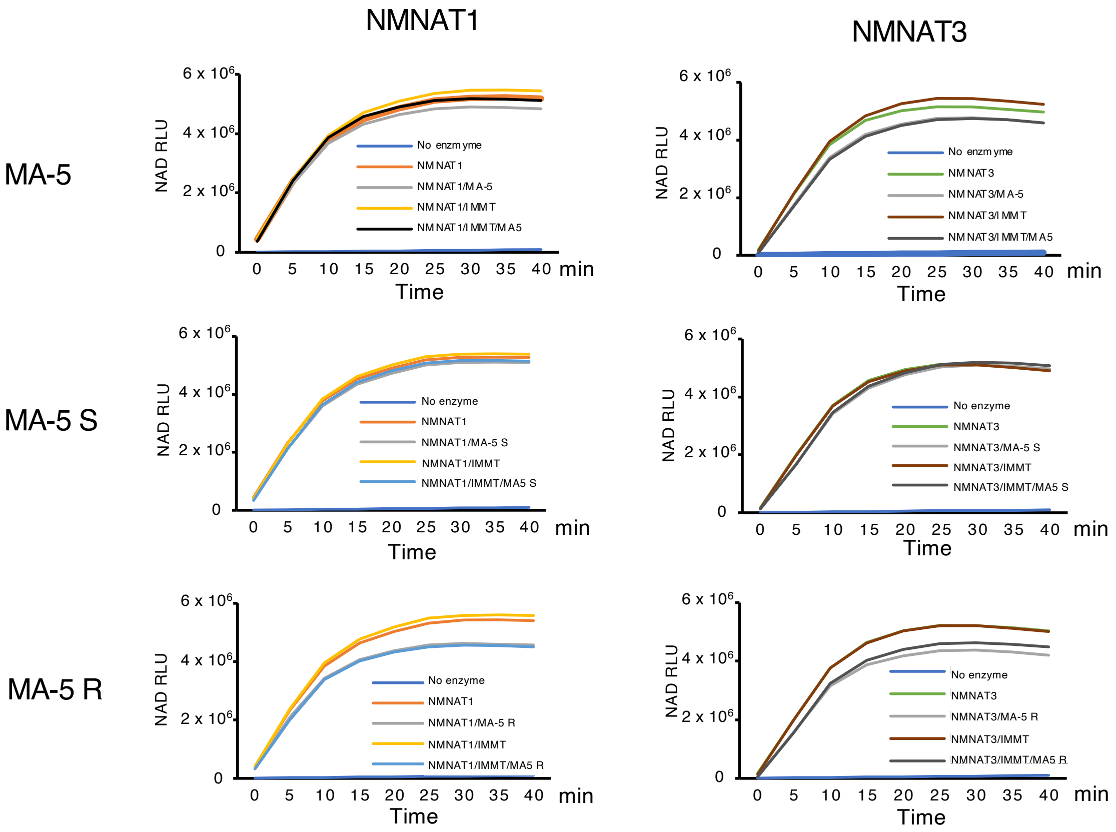

### Extended Data Figure 4

a

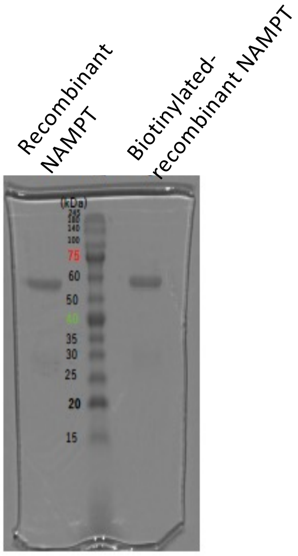


b

#### Binding assay for NAMPT

①ビオチン化NAMPT/100mM S体
(PBS, ビオシチン50μM, NAMPT0.04μg/200μL)

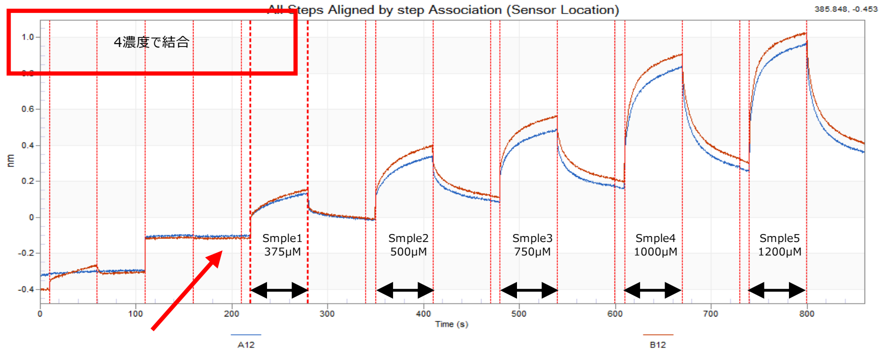


②ビオチン化NAMPT/100mM R体
(PBS, ビオシチン50μM, NAMPT0.04μg/200μL)

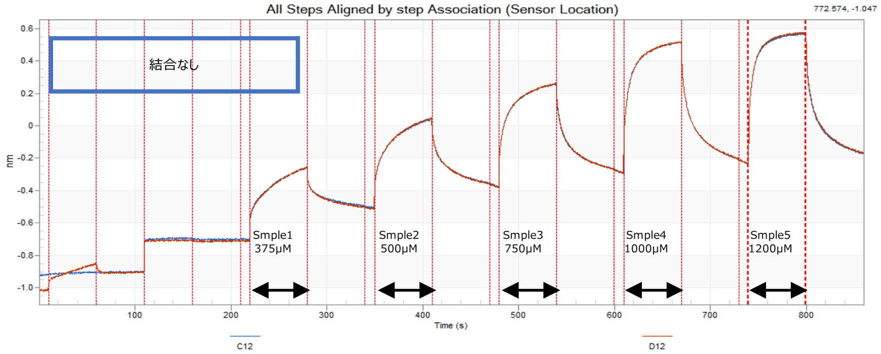

### Extended Data Figure 5

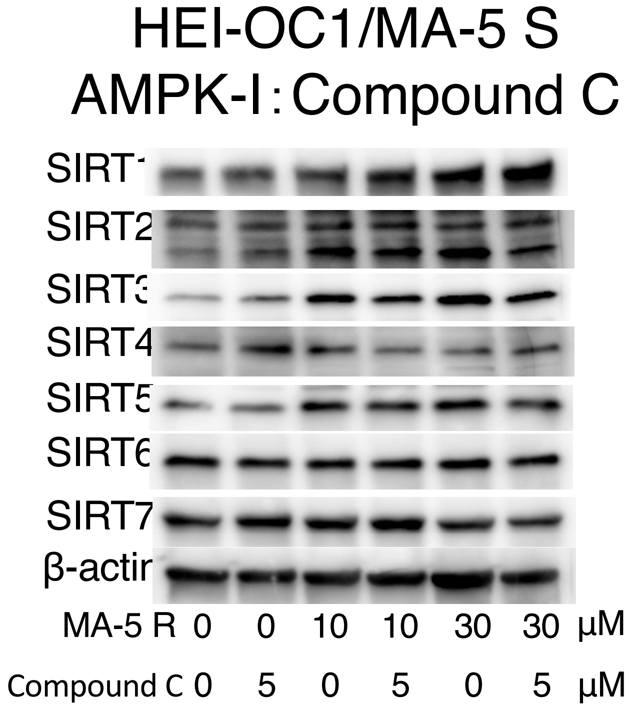

### Extended Data Figure 6

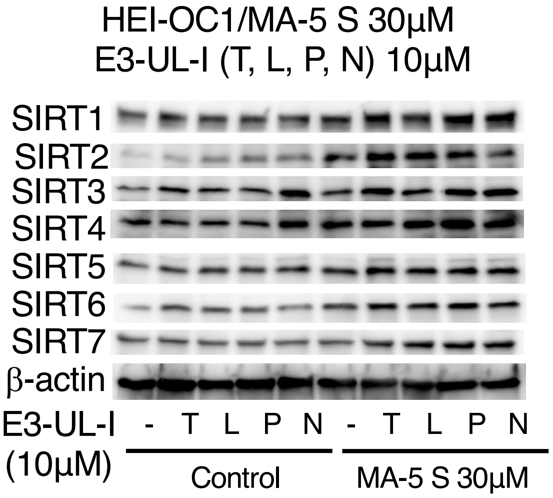


T : Thalidomide 10μM, L: Lenalidomide 10μM,
P : Pomalidomide 10μM, N : NSC 66811 10μM
E3-UL-I : E3 Ubiquitin ligase inhibitor

### Extended Data Figure 7

a

Predicted binding mode of S-enantiomer to DNA-PKcs using MOE.

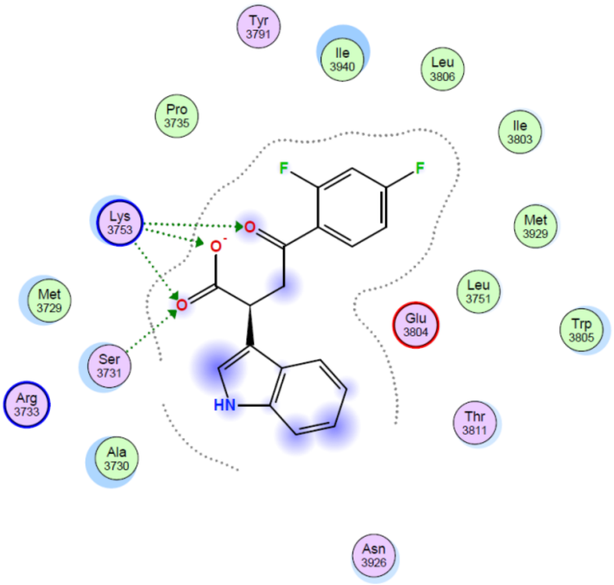


Predicted complex structure of S-enantiomer to DNA-PKcs using MOE.

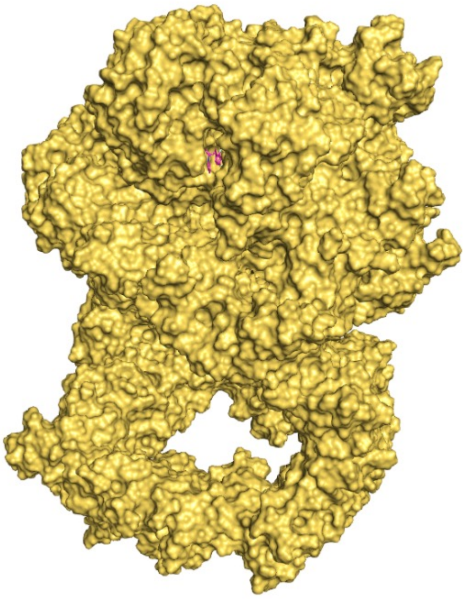

b

Binding mode of M3814 to DNA-PKcs (PDB ID: 7OTY).

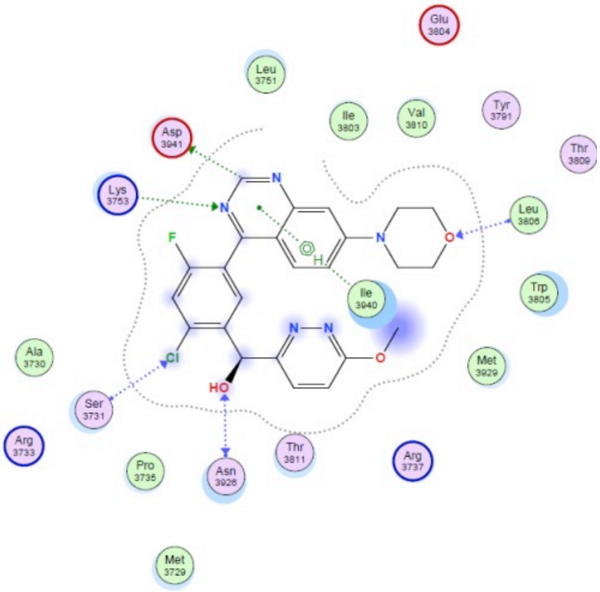


Complex structure of M3814 to DNA-PKcs (PDB ID: 7OTY).

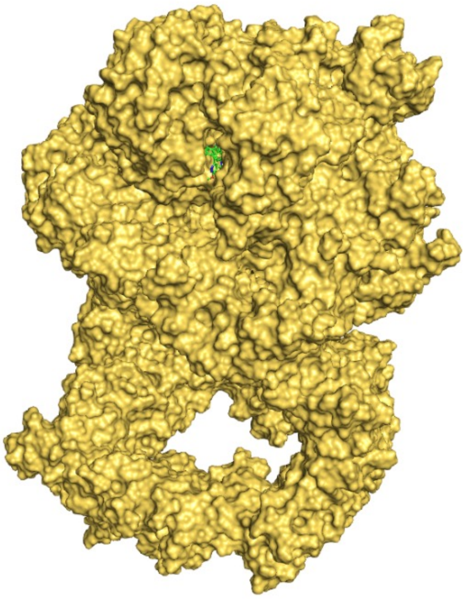

### Extended Data Figure 8

a

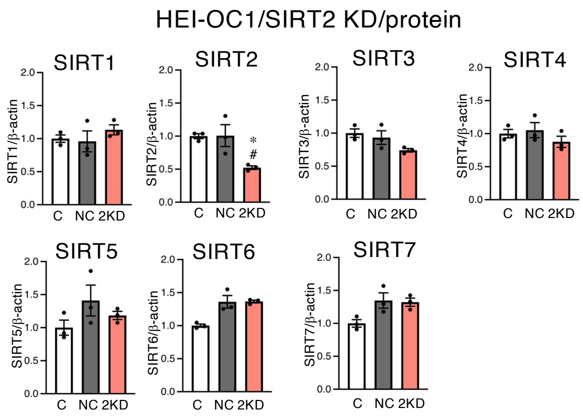


b

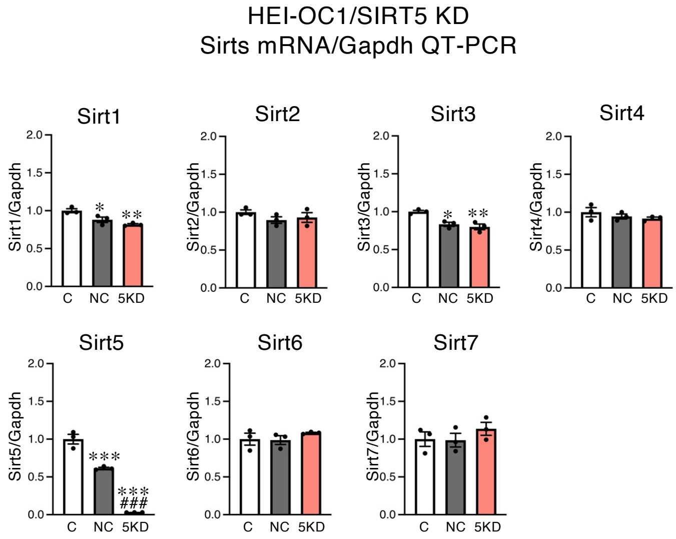


c

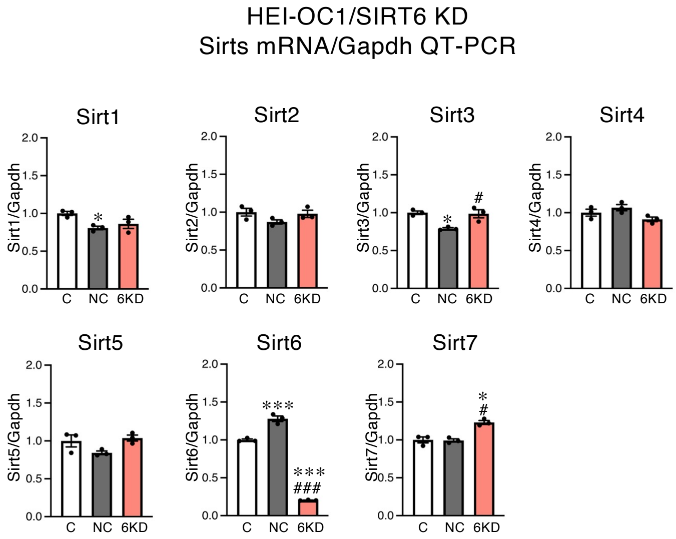

### Extended Data Table 1

TaqMan Primers were purchased from Thermo Fisher Scientific (**Table S1**).

| PCR-Primers | Source | Identifer |
| --- | --- | --- |
| murine Sirt1 | Applied Bio systems/Thermo Fisher Scientific | Mm01168521_m1, Cat # 4331182 |
| murine Sirt2 | Applied Bio systems/Thermo Fisher Scientific | Mm01149204_m1, Cat # 4331182 |
| murine Sirt3 | Applied Bio systems/Thermo Fisher Scientific | Mm00452131_m1, Cat # 4331182 |
| murine Sirt4 | Applied Bio systems/Thermo Fisher Scientific | Mm01201915_m1, Cat # 4331182 |
| murine Sirt5 | Applied Bio systems/Thermo Fisher Scientific | Mm00663723_m1, Cat # 4331182 |
| murine Sirt6 | Applied Bio systems/Thermo Fisher Scientific | Mm01149042_m1, Cat # 4331182 |
| murine Sirt7 | Applied Bio systems/Thermo Fisher Scientific | Mm01248607_m1, Cat # 4331182 |
| murine Prkdc(DNA-PKcs) | Applied Bio systems/Thermo Fisher Scientific | Mm01342967_m1, Cat # 4331182 |
| murine Atm | Applied Bio systems/Thermo Fisher Scientific | Mm00554903_m, Cat # 4331182 |
| murine Gapdh | Applied Bio systems/Thermo Fisher Scientific | Mm99999915_g1 Cat # 4331182 |
| murine Actb(beta-actin) | Applied Bio systems/Thermo Fisher Scientific | Mm02619580_g1, Cat # 4331182 |

siRNAs were purchased from Applied Bio systems/Thermo Fisher Scientific (**Table S2**).

| siRNA | Source | Identifer |
| --- | --- | --- |
| murine Prkdc(DNA-PKcs) , Silencer® Select Pre-Designed siRNA | Thermo Fisher Scientific | s201831, Cat #. 4390771 |
| murine Atm, Silencer® Select Pre-Designed siRNA | Thermo Fisher Scientific | s62692, Cat #. 4390771 |
| murine Sirt2, Silencer® Select Pre-Designed siRNA | Thermo Fisher Scientific | s82217, Cat #. 4390771 |
| murine Sirt5, Silencer® Select Pre-Designed siRNA | Thermo Fisher Scientific | s86590, Cat #. 4390771 |
| murine Sirt6, Silencer® Select Pre-Designed siRNA | Thermo Fisher Scientific | s78392, Cat #. 4390771 |
| Silencer™ Select Negative Control No. 1 siRNA | Invitrogen | Cat #. 4390843 |
| Silencer™ Select Negative Control No. 2 siRNA | Invitrogen | Cat #. 4390846 |
